# Cardiomyocyte-specific loss of *Smyd5* leads to a robust activation of inflammatory signaling and heart failure in mice

**DOI:** 10.64898/2026.09.02.748923

**Authors:** Ryan Bia, Samuel Hickenlooper, Mickey R. Miller, Caiyi C. Li, Emilee Horiuchi, Anna Bakhtina, Li Wang, Steven Valdez, Kathryn Davis, Alexa Anderson, June García-Llana, Ludovica Farese, Sean O’Very, Nicholas Santa Ana, Anna Jacobsen, Kyra Dellerman, Clint Gwynn, Jessica Durrant, Joseph R Visker, Christos P. Kyriakopoulos, Konstantinos Sideris, Stavros G. Drakos, Marta W. Szulik, Edward B. Thorp, Stephen T. Smale, Sarah Franklin

**Author notes:** Denoted authors contributed equally to this project (Ryan Bia, Samuel Hickenlooper, Mickey R. Miller, Sarah Franklin). Correspondence: Sarah Franklin Department of Internal Medicine Nora Eccles Harrison Cardiovascular Research and Training Institute University of Utah Salt Lake City, UT.

## Abstract

**Background:** Cardiomyocytes respond to stress by undergoing hypertrophic growth driven by dynamic changes in gene expression. Epigenetic mechanisms, including histone methylation, play critical roles in regulating these transcriptional programs, yet the enzymes controlling these modifications during cardiac disease remain largely unknown. The SMYD family of histone methyltransferases regulates gene expression in multiple biological contexts, but the function of SMYD5 in the mammalian heart has never been investigated.

**Methods:** SMYD5 expression was assessed in human heart failure samples and in a mouse model of cardiac hypertrophy. To define its functional role in vivo, we generated inducible cardiomyocyte-specific *Smyd5* knockout mice and characterized their cardiac phenotype using molecular, histological, and functional analyses. Chromatin immunoprecipitation-quantitative PCR (ChIP-qPCR) was performed to examine histone H4 lysine 20 trimethylation (H4K20me3) at the *Il-6* promoter.

**Results:** SMYD5 expression was altered in diseased human and mouse hearts. Under basal conditions, cardiomyocyte-specific deletion of *Smyd5* resulted in baseline structural cardiac remodeling and transcriptional signatures characteristic of pathological stress. *Smyd5*-deficient hearts exhibited marked inflammatory activation resembling a cytokine storm with immune cell infiltration and heart failure. Notably, *Smyd5* knockout mice displayed a 100-fold increase in *Il-6* expression, accompanied by a global reduction in H4K20me3. ChIP-qPCR analysis of the *Il-6* promoter, together with loss- and gain-of-function analysis of SMYD5, supports a direct epigenetic role of SMYD5 in regulating *Il-6* expression through H4K20me3 in cardiomyocytes.

**Conclusions:** SMYD5 is a previously unrecognized epigenetic regulator of cardiac homeostasis that restrains inflammatory signaling in cardiomyocytes under normal conditions. Loss of *Smyd5* disrupts H4K20me3, leading to derepression of *Il-6* in cardiomyocytes and a robust inflammatory response characterized by immune cell recruitment and fibrosis, accompanied by rapid progression of cardiac remodeling and heart failure. These findings identify SMYD5 as a critical regulator of intrinsic cardiomyocyte inflammatory signaling and reveal a novel chromatin-based mechanism contributing to inflammatory cardiomyopathies.

**NOVELTY AND SIGNIFICANCE:** *What Is Known?:*

- Elevated levels of pro-inflammatory cytokines such as IL-6 are strongly associated with adverse cardiac remodeling and poor outcomes in heart failure.
- Histone methylation is a key epigenetic mechanism regulating gene expression in the heart, but the role of histone H4K20 trimethylation and its regulatory enzymes in cardiomyocytes remains poorly understood.
- The histone methyltransferase SMYD5 regulates gene expression and inflammatory pathways in several non-cardiac cell types, but its role in the mammalian heart has not been examined.

*What New Information Does This Article Contribute?:*

- This study provides the first in vivo characterization of SMYD5 in the heart and identifies it as a critical epigenetic regulator of cardiomyocyte homeostasis.
- Cardiomyocyte-specific loss of *Smyd5* triggers pathological hypertrophy, elevated inflammatory signaling, immune cell infiltration, fibrosis, and heart failure.
- SMYD5 directly represses *Il-6* expression in cardiomyocytes through histone H4K20 trimethylation at the *Il-6* promoter, revealing a previously unrecognized epigenetic mechanism controlling intrinsic cardiomyocyte-driven inflammation. Epigenetic mechanisms that regulate inflammatory signaling in cardiomyocytes remain largely unknown. Here, we identify the histone methyltransferase SMYD5 as a critical regulator of cardiac homeostasis and intrinsic inflammatory signaling. Using an inducible cardiomyocyte-specific *Smyd5* knockout mouse model, we demonstrate that loss of *Smyd5* induces rapid progression to heart failure accompanied by robust inflammatory activation, including a ∼100-fold increase in *Il-6* expression, inflammatory immune cell infiltration, fibrosis, and severe cardiac dysfunction. Mechanistically, SMYD5 directly regulates *Il-6* expression by catalyzing histone H4 lysine 20 trimethylation at the *Il-6* promoter, thereby restraining pro-inflammatory gene expression in cardiomyocytes. Ablation of *Smyd5* markedly reduces global H4K20 trimethylation, and results in dramatic upregulation of *Il-6* and downstream cytokine signaling pathways, producing a phenotype resembling cytokine storm-like inflammatory cardiomyopathy. These findings establish SMYD5 as the first epigenetic regulator shown to suppress intrinsic cardiomyocyte inflammatory signaling and uncover a novel chromatin-based mechanism controlling cytokine production in the heart. Targeting SMYD5-dependent pathways therefore represents a new strategy for limiting maladaptive inflammation in heart failure and inflammatory cardiomyopathies.

## INTRODUCTION

Inflammation is typically a protective response to infection or tissue injury, however, prolonged or non-resolved inflammation can also damage tissue and is considered a key component of the pathogenesis of cardiac diseases, including heart failure and adverse ventricular remodeling^1^. Indeed, pro-inflammatory cytokines, including Tumor Necrosis Factor alpha (TNFα) and Interleukin-6 (IL-6), which are released from cardiomyocytes as part of the initial inflammatory response, are elevated in heart failure patients. Circulating levels of these two proteins have been shown to independently predict mortality^2,3^. It is also of significant interest that one common feature between heart failure patients with reduced or preserved ejection fraction is the correlation between elevated serum pro-inflammatory cytokines and adverse clinical outcomes^4–7^.

While cardiomyocytes can initiate the inflammatory cascade by secreting these pro-inflammatory cytokines, this process is further propagated by other non-myocyte cells, including fibroblasts, neutrophils, and macrophages. Fibrosis is the final step in the inflammatory process, which results from accelerated collagen deposition and is a leading cause of heart failure in patients after myocardial infarction. Current therapies to combat prolonged inflammation have included inactivating cytokines, reducing the fibroblast response, or reversing fibrosis, however, these have been largely ineffective^8–10^. Therefore, controlling the initial cytokine production from cardiomyocytes may be one key strategy for preventing or minimizing tissue damage. Indeed, studies in animals aimed at blocking IL-6 production (via *Il-6* knockout mice) show that deletion of *Il-6* attenuates pressure overload-induced hypertrophy and cardiac dysfunction^11^, and ameliorates myocardial remodeling after myocardial infarction^12^, suggesting that targeting this pathway therapeutically could be beneficial. While our understanding of the molecular processes regulating cytokine production and release from cardiomyocytes has improved in recent years, we still know very little regarding the underlying epigenetic mechanisms driving these transcriptional changes and how they contribute to disease.

On a molecular level, it has been well established that many cardiac pathologies result from dynamic changes in gene expression and, conversely, that modulating key epigenetic factors (e.g., BRG1^13^, BET^14^, HDACs^15^) have the capacity to prevent or abrogate injury or pathological remodeling. However, we are only beginning to identify the epigenetic mechanisms in the heart that regulate these transcriptional changes. The methylation of lysine residues on histone proteins has been shown to be a key driver of gene expression and has been correlated with gene activation or silencing, depending on the residue modified, with additional complexity arising from the ability of each lysine to be mono-, di-, or tri-methylated. Much research has focused on understanding the functional significance of methylation on histone H3 lysine 4 (K4) and lysine 9 (K9), marks of gene activation and silencing, respectively. However, much less is known regarding methylation on other lysine residues, including histone H4 lysine 20 (K20). Methylation of histone H4K20 has been implicated in gene activation, heterochromatin formation, cell cycle, DNA repair, and gene silencing in non-cardiac cells, depending on the degree of methylation (mono-, di-, or tri-), however, this modification has not been well studied in cardiomyocytes. In the heart, only two recent studies have examined global levels of H4K20 in cell and rodent models and showed decreased trimethylation of histone H4K20 during cardiac hypertrophy and heart failure, although the specific genes regulated by these changes are largely unknown^16,17^. In addition, the enzymes that add or remove these methylation marks have only begun to be identified but have recently been linked to the development of congenital heart disease, cardiac hypertrophy, and heart failure^18,19^.

The SMYD family is a unique family of methyltransferases that have been shown to regulate gene expression through the selective addition of methyl groups on histone proteins. Since the discovery of this family SMYD1, 2, and 3 have been studied for their involvement in cell growth and differentiation in various cell types during development and disease^20,21,22^. However, the remaining two family members, SMYD4 and SMYD5, have remained largely uncharacterized.

To date, SMYD5 has been examined in seventeen primary research articles using cultured cells (embryonic stem cells^23,24^, macrophages^25^, and intestinal epithelial cells^26^), zebrafish embryos^27^, or a mouse model of rheumatoid arthritis^28^, although no study has examined its role in the mammalian heart (Table S1). Specifically, these previous studies have shown that SMYD5 influences cell renewal^23^ and differentiation^24^ in embryonic stem cells. In intestinal epithelial cells, SMYD5 methylates PGC-1α, thereby regulating the protein’s stability and degradation, which affects mitochondrial function in inflammatory bowel disease^26^. In cultured macrophage cells (RAW 264.7), histone methylation by SMYD5 was shown to play a significant role in inflammation by regulating the expression of Toll-Like Receptor 4 (TLR4) target genes, including *Il-1a, Tnfα*, *Cxcl10*, and *Ccl4*^25^. Additionally, in developing zebrafish embryos^27^, morpholino knockdown of *smyd5* showed normal gross morphological development, including heart and skeletal muscle, concomitant with increased expression of both primitive and definitive hematopoietic markers, suggesting Smyd5 influences hematopoiesis^27^. In the knee joints of mice, SMYD5 regulated synovial fibroblast homeostasis and the pathogenesis of rheumatoid arthritis. This study demonstrated reciprocal regulation of *Il-6* expression in fibroblast-like synoviocytes, where SMYD5 overexpression enhanced *Il-6* production, whereas *Smyd5* knockdown suppressed it, contributing to alleviated joint inflammation and arthritis severity in the collagen-induced arthritis mouse model^28^. Mechanistically, SMYD5 has been shown to regulate gene expression through site-specific methylation of histone H4K20, H3K36, and H3K37 at gene promoters^29–32^.

However, no study has examined the role of SMYD5 in the adult heart or evaluated its capacity to regulate cardiomyocyte gene expression, pathological remodeling, or cardiac dysfunction. To address this gap, we first assessed SMYD5 expression in the myocardium of human heart failure patients and in mouse model of cardiac disease. We then generated and comprehensively characterized inducible, cardiomyocyte-specific *Smyd5* knockout mice to define its functional role in vivo. In parallel, we employed complementary in vitro approaches utilizing targeted gain- and loss-of-function to elucidate the molecular basis for SMYD5-mediated regulation of intrinsic cardiomyocyte-driven inflammatory signaling. Together, this work constitutes the first study of SMYD5 in cardiac biology and establishes SMYD5 as a novel epigenetic regulator of inflammatory cardiomyopathy and cardiac homeostasis.

## METHODS

### Human Heart Tissue Acquisition

The study was conducted in accordance with the Declaration of Helsinki and approved by the Institutional Review Board of the University of Utah. All samples were collected with written informed consent. Human heart tissue samples were acquired as previously published^33^. We prospectively enrolled patients (age≥18-years) in institutions comprising the Utah Transplantation Affiliated Hospitals (U.T.A.H.) Cardiac Transplant Program (i.e. University of Utah Health Science Center, Intermountain Medical Center, and the Veterans Administration Salt Lake City Health Care System), who had clinical characteristics consistent with dilated cardiomyopathy and chronic advanced heart failure, who required circulatory support with continuous flow Left Ventricular Assist Device (LVAD) as a bridge to transplantation or lifetime destination therapy. Patients who required LVAD support due to acute heart failure (acute myocardial infarction, acute myocarditis, post-cardiotomy cardiogenic shock, etc.) were prospectively excluded. LVAD patients underwent serial echocardiograms monthly for the first three months, then at four and a half and six months. They were categorized as either responders or non-responders using left ventricular ejection fraction (LVEF) and left ventricular end-diastolic diameter (LVEDD) measurements during diminished LVAD support “turn-down” echocardiograms. Responders were defined as patients with a final LVEF >40% and LVEDD ≤5.9cm, whereas non-responders were defined as patients with a final LVEF <35% and with <50% relative improvement in LVEF regardless of the final LVEDD. For heart failure patients, their clinical demographics, echocardiographic parameters, protein biomarkers, and other clinical data were prospectively collected and entered in our program’s research electronic data capture system (REDCap) and were previously published^33^. Myocardial tissue was prospectively collected from the LV apical core at the time of LVAD implantation and was snap frozen before storing it at -80°C, as described before^34,35^. Myocardial tissue from donor hearts, not allocated for heart transplantation due to non-cardiac reasons (size, infection, etc.), was used as non-failing controls, and LV apical tissue was harvested and processed the same way as heart failure patients. It is important to note that some myocardial tissue samples and their respective ejection fractions from patients with HF and donors in the original study were used for both RNA sequencing and proteomics experiments, whereas others were used in just one application; therefore, the sum of ejection fraction measurements includes all cardiac tissue samples analyzed previously. The RNA-Seq data shown in Figure 1F was previously generated as part of a large dataset by Drakos et al.^33^ and described in that study as follows: “differential gene expression was determined using DESeq2 package, which applies the Benjamini-Hochberg method for multiple testing correction. The criteria for differential expression included: adjusted *p*-value <0.05, absolute log_2_fold change >0.585 (equivalent to linear fold change >1.5), and a normalized base mean count of 30 (to remove any significant genes with very low expression”^33^. *SMYD5* did not meet the fold change criteria applied to the published portion of the dataset. Therefore, we have used the unpublished portion of this data set to extract log2 values for *SMYD5* and perform an analysis of transcript abundance between failing (responders and non-responders) and healthy heart samples. The statistical analysis was performed using a two-way ANOVA with a post hoc Tukey’s test for multiple comparisons. For a more targeted analysis of *SMYD5* abundance using RT-qPCR, we utilized cardiac tissue from a separate cohort of patients. The clinical characteristics of this additional study population, including clinical demographic data, heart failure etiology, and echocardiographic parameters, have been described previously^36^.

**Figure 1:**
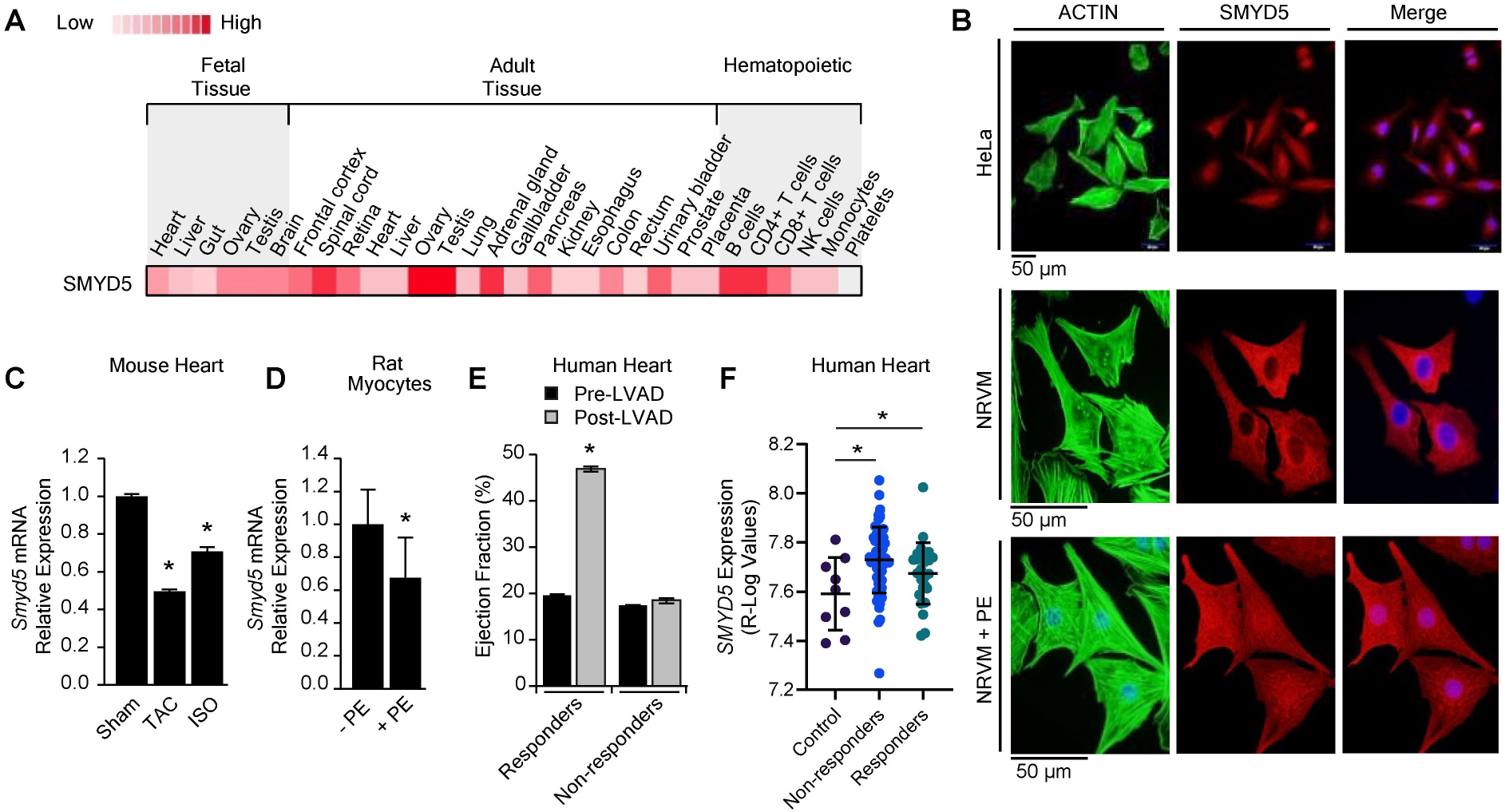
SMYD5 expression is decreased in the diseased heart. **A**, SMYD5 protein expression was assessed in tissue and cell types generated from the Human Proteome Map portal. **B**, Immunostaining of HeLa and NRVM cells (with or without phenylephrine (PE) treatment to induce hypertrophy) using phalloidin (green), anti-SMYD5 antibody (red), and DAPI (blue) to determine subcellular localization of SMYD5. Evaluated 40 cells per group in three independent experiments. **C**, Mice undergoing TAC or ISO treatment via mini-osmotic pump implantation showed a decrease in *Smyd5* expression as measured via RT-qPCR. *n*=3. A one-way ANOVA was used for statistical analysis. Asterisk * indicates *p*-value <0.01. **D**, In NRVMs treated with PE, *Smyd5* gene expression was downregulated. *n*=3. A two-tailed unpaired Student’s *t*-test was used for statistical analysis. Asterisk * indicates *p*-value <0.05. **E,** Comparison of ejection fraction in human heart failure patients who underwent LVAD implantation. *n*=25 (Responders), *n*=59 (Non-responders). A two-way ANOVA with the Tukey-Kramer test for multiple comparisons was used for statistical analysis. Asterisk * indicates *p*-value <0.01. **F,** *SMYD5* expression was increased in heart failure patients regardless of their capacity for functional recovery via LVAD unloading as determined by bulk RNA-Seq. *n*=9 (Control), *n*=59 (Non-responders), *n*=25 (Responders). A one-way ANOVA was used for statistical analysis. Asterisk * indicates *p*-value <0.01. Data are expressed as the mean ± SEM (**C–F**).

### Inducible Cardiac-Specific Smyd5 Gene Knockout Mouse

All protocols involving animals conform to the NIH Guide for the Care and Use of Laboratory Animals and were approved by the Institutional Animal Care and Use Committee of the University of Utah. All efforts were made to minimize pain and distress during procedures and when harvesting cardiac tissue under anesthesia. In this study, we used both male and female C57Bl/6 mice. To develop the targeting construct to generate Smyd5^flox/flox^ mice, a small fragment containing exon 2 was cloned in between two *loxP* sites. Two 2-3kb fragments flanking exon 2 were PCR-amplified and cloned outside of the *loxP* sites as homologous arms. After in vivo CRE-mediated recombination, the floxed allele and the null allele were screened by PCR. Cardiac-specific inducible iSmyd5 knockout (KO) mice were generated by crossing Smyd5^flox/flox^ mice with *α-MHC-MerCreMer* mice purchased from Jackson Laboratory (Cat. no. 005657). These mice are referred to as iSmyd5 throughout the manuscript. To induce recombination, adult mice between the ages of 8-13 weeks were fed a diet containing tamoxifen (TMX), 0.4 mg per gram of chow diet (Harlan, Cat no. TD.130859), and were monitored carefully to ensure ingestion. Control mice utilized in these experiments are defined as age-matched littermates that carry the floxed *Smyd5* gene but lack the *Cre*-recombinase transgene or carry the *Cre-*recombinase transgene and lack floxed *Smyd5* (both are incapable of deleting exon 2), and are also fed a tamoxifen-chow diet for the same duration of the experiment. We have also examined an additional control where floxed, *Cre*-positive mice are maintained on normal chow and have compared these results to mice fed TMX chow and see no effect of tamoxifen on basal cardiac function in this strain. Knockout of *Smyd5* in the heart was confirmed at both the protein and RNA levels via immunoblotting and RT-qPCR, respectively. Skeletal muscle tissue was also examined via immunoblot and RT-qPCR to ensure specific knockdown of *Smyd5* only in the heart.

### Transverse aortic constriction (TAC) surgery

The mouse model of transverse aortic banding-induced cardiac hypertrophy was performed as described previously^37,38^. Briefly, the 8-12 week-old mice were randomly assigned to two groups: TAC (treated group, *n*=3) or Sham (control group, *n*=3). Mice were anesthetized with 1.5-3.0% isoflurane, intubated, and ventilated with 1.5-2.0% isoflurane in 98% O_2_/2% CO_2_. After shaving the hair from the mouse, the chest was entered from the left side via the third intercostal space. The aorta was identified at the T8 region, and a venous vascular clamp outfitted with a band of silastic tubing at the distal edge of one of the clamps was placed around the vessel. The internal diameter of the resulting modified clamp measured 27G. Once the aorta was clamped, distal blood flow was measured and quantified using a flow probe. The chest was then closed using a 6-0 Prolene suture, during which negative pressure in the thorax was reestablished by removing air with a PE-50 chest tube attached to a syringe. Sham-operated mice underwent the same procedure without aortic clamping.

### ISO-induced model of cardiac hypertrophy in mice

The murine model of isoproterenol-induced cardiac hypertrophy via mini-osmotic pump was performed as described previously^39^. Briefly, 10-15 week-old mice were randomly assigned to two groups: ISO (treated group, *n*=11) or Sham (control group, *n*=9). Mini-osmotic pumps filled with isoproterenol (ISO) or saline were surgically implanted into mice, which administered 30 mg/kg/day for 4 weeks with the Alzet model 2006, releasing 0.15μl per hour.

### Echocardiography

Echocardiography (ECHO) was performed on a Vevo 2100 (Visual Sonics) to determine cardiac parameters in live mice, as described previously^37,38^. Parameters measured include ventricular function (ejection fraction [EF], fractional shortening [FS], stroke volume [SV], left ventricular end-diastolic diameter [LVEDD], and left ventricular end-systolic diameter [LVESD]).

### Histology

At the time of sacrifice, iSmyd5 mice were anesthetized, and whole hearts were rapidly excised from the animals and soaked in 1M KCl prepared in 1× PBS to arrest them in diastole. The hearts were fixed in 4% paraformaldehyde and then embedded in paraffin. Hearts were then sectioned at 4μm and placed on slides. Hematoxylin and Eosin Staining (H&E) and Masson’s Trichrome (MT) staining were performed by the Huntsman Cancer Institute Biorepository and Molecular Pathology Research Histology Core. Cardiac tissue sections were then incubated with Texas red-X-conjugated wheat germ agglutinin (Invitrogen, 1:100) for 90 minutes and assessed for cell size. Tissue sections were visualized and imaged on an Olympus BX51WI microscope equipped with a DP73 color camera, and image analysis was performed with CellSens Standard software (Olympus).

To quantify the area of immune infiltration and collagen deposition, MT and H&E slides were evaluated by an ACVP board-certified veterinary pathologist. One MT image per heart (*n* = 4/group) was imported to ImageJ software for measurement of infiltration and collagen. H&E-stained slides (*n* = 4/group) containing multiple sections of the heart per animal were also scored according to the extent of infiltrated area: 0 = absent, 0.5 = very minimal (<1% affected), 1 = minimal (1-10% affected), 2 = mild (10–20% affected), 3 = moderate (21–30% affected), 4 = marked (31–40% affected), 5 = severe (> 40% affected) ^40^. The scores for each section were averaged for individual animals. In addition, cardiomyocyte diameter was measured in 100 cardiomyocytes (60 in the LV free wall, 40 in the septum) in randomly chosen fields for each sample using ImageJ software. The mean diameter for the free wall and septum was determined, and the standard deviation was calculated (to assess variability).

### Adult Cardiomyocyte Isolation via Langendorff Perfusion Method

Isolation of cardiomyocytes and fibroblasts from cardiac tissue harvested from iSmyd5 KO and control (Ctrl) mice was carried out from week one and week two time points after TMX administration, utilizing the Langendorff retrograde perfusion technique, as described previously^41^. Briefly, mice were administered heparin via intraperitoneal injection, followed one minute later by a pentobarbital injection. Oxygen was administered to the animal, and surgical removal of the heart was carefully performed once the animal was fully anesthetized. After removal, the heart was then tied to a cannula via the aorta with two sutures. The heart was then washed with cold Ca^2+^-free Tyrode’s solution by attaching the cannula with the heart to a syringe. The cannula with the heart attached was then moved to the perfusion system, where the heart was then washed with warm Ca^2+^-free Tyrode’s solution to remove any blood remaining in the chambers/vessels. Enzyme solution containing Collagenase Type 2 (Worthington Biochemical Corporation) and Protease 14 (Sigma Aldrich) was then added to the perfusion system for 18 minutes, followed by an enzyme stop buffer to stop the enzymatic reaction. The ventricles of the heart were then isolated from the atrium and finely minced in the stop buffer. The cells were mechanically dissociated by triturating with a serological pipette and strained into a sterile 15 mL centrifuge tube. Cardiomyocytes and fibroblasts were then separated by multiple gravity settling steps for purification and removal of dead cells. Cardiomyocytes were then examined under the microscope to determine cardiomyocyte and fibroblast viability. Quantification of cell-specific markers *Myh6*, *Col1a1*, and *Vim* via RT-qPCR, as detailed below, was carried out to confirm cell type enrichment (primers listed in Table S2). These cell populations were then utilized to quantify *Smyd5* expression.

### Neonatal Cardiomyocytes and Primary Cardiac Fibroblasts Isolation

Neonatal rat ventricular myocytes (NRVMs) were extracted as previously published ^20^ from 24-hour old pups of embryonic stage 21 Sprague Dawley rats using the manufacturer’s protocol (Worthington Biochemical Corporation) with modifications to separate fibroblasts from cardiomyocytes. Both NRVMs and primary cardiac fibroblasts were plated in 10% fetal bovine serum (FBS), 1% penicillin-streptomycin (P/S), 1% non-essential amino acid DMEM media for 24 hours, after which the media was switched to serum-free media supplemented with 1% ITS and 1% P/S for further culturing of NRVMs. Primary cardiac fibroblasts were cultured in 10% fetal bovine serum (FBS), 1% penicillin-streptomycin (P/S), 1% non-essential amino acid DMEM media. For imaging, NRVMs were plated at 10% confluency on glass coverslips pre-treated with 50 μg/ml laminin (BD Biosciences). After 24 hours the cells were fixed by incubation with 4% paraformaldehyde solution (Santa Cruz) for immunocytochemical staining. In addition, separate NRVMs and primary cardiac fibroblasts were cultured and subjected to *Smyd5* knockdown or adenoviral-mediated SMYD5 expression as detailed below in the presence or absence of phenylephrine (PE, Sigma Aldrich; 50μM for 48 hours).

### Cell Culture

HeLa or H9c2 cells were cultured in 10% FBS with 1% P/S, changing media as needed, until they were 70% confluent, at which point cells were split. Passage numbers were kept under 10.

### Adenovirus Expression

NRVMs, H9c2 or HeLa cells were transduced with adenovirus expressing FLAG-tagged SMYD5 (Vector Biolabs) at the multiplicity of infection (MOI) of 25. All cells were cultured for 24 hours in the presence or absence of PE, as we have previously published ^20^. Control cells were transduced with an empty control virus (Ad-Null) and served as a negative control in these experiments.

### siRNA Knockdown

To induce *Smyd5* knockdown, NRVMs or cardiac fibroblast cells, were transfected with si-*Smyd5*-RNA (Qiagen) as previously published, and efficiency of knockdown confirmed^21^. Control cells were transfected with scrambled siRNA (Qiagen).

### Immunocytochemical staining

NRVMs and HeLa cells were fixed using a 4% paraformaldehyde solution on glass coverslips and permeabilized by incubation with 0.25% Triton X-100 (Sigma). Blocking was carried out by incubating the cells in 10% bovine serum albumin. The cells were then incubated with anti-Myc antibody (Abcam, ab9106) followed by incubation with Alexa Fluor® 568 goat anti-rabbit secondary antibody (Thermo Fisher Scientific, A11011) as well as Alexa Fluor® 488 phalloidin (Thermo Fisher Scientific, A12379). The coverslips were then mounted on glass slides using Prolong® Gold antifade reagent with DAPI (Thermo Fisher Scientific, P36935) and allowed to adhere overnight. Cells were visualized and imaged on an Olympus BX51WI microscope with CellSens Standard software (Olympus).

### Electrophoresis and immunoblotting

Proteins were separated by standard SDS-PAGE using Laemmli buffer. For immunoblotting, proteins were transferred to nitrocellulose, membranes blocked with 10% powdered milk reconstituted in 1× TBST, and protein signals detected by enzyme-linked chemiluminescence (Thermo Fisher Scientific). Ponceau staining of membranes was performed before blocking and used to confirm transfer and protein loading. Antibodies used in this study are as follows: Histone H3 (Abcam, ab1791); Histone H3-trimethylated-K4 (Abcam, ab8580); Histone H3-trimethylated-K9 (Abcam, ab8898); Histone H4 (Abcam, ab10158); Histone H4-trimethylated-K20 (Abcam, ab9053); β-Tubulin (Abcam, ab6046); FLAG (Sigma, F1804); SMYD1 (Abcam, ab32482); SMYD2 (Abcam, ab108217); SMYD3 (Abcam, ab187149); SMYD4 (Thermo Fisher Scientific, PA5-96631); SMYD5 (Abcam, ab81419); goat anti-rabbit IgG (Abcam, ab97051).

### Quantitative Real Time PCR Analysis

Total RNA was isolated using TRIzol (Invitrogen) according to the manufacturer’s protocol. Total RNA was transcribed using SuperScript III First-Strand Synthesis system for RT-qPCR (Invitrogen) according to the manufacturer’s protocol to produce cDNA. The cDNA transcripts were amplified on the Bio-Rad CFX Connect Real-Time System using SsoAdvanced Universal SYBR Green Supermix (Bio-Rad). Gene expression analysis was performed using the 2^-ΔΔCt^ method, using either *β-Tubulin*, *α*-*Actin*, or *Gapdh* as a control housekeeping gene. Oligonucleotide primers used in this study were previously published and designed using NCBI Primer-BLAST^42^ and obtained from Thermo Fisher Scientific. A list of primers used in this study are provided in Table S2.

### Chromatin Immunoprecipitation

Chromatin was isolated from NRVMs that had been infected with FLAG-tagged SMYD5 (MOI 25 for 24 hours). An empty vector was used as a negative control in this experiment. Chromatin immunoprecipitation was performed using a commercially available ChIP-IT High Sensitivity kit (Active Motif, 53040) according to the manufacturer’s instructions. Chromatin-bound proteins were immunoprecipitated using anti-H4K20me3 (Abcam, ab9053) and anti-FLAG (Sigma-Aldrich F1804). Immunoprecipitated DNA was analyzed by RT-qPCR using primer sets that amplified the promoter region of *Il-6*. *Tbp* and an intergenic region as negative controls. qPCR was performed in duplicate with equal immunoprecipitated samples and input. Values were normalized to input measurements, and enrichment was calculated using the ΔΔCt method.

### In-solution Digestion of Mouse Heart Tissue for Mass Spectrometry

Heart tissue was placed in a buffer containing 4% SDS, 100 mM Tris (pH 7.6), and protease inhibitors. Samples were sonicated with three 10-second pulses using a sonic dismembrator (Fisher) at 25% amplitude. The crude extract was clarified by centrifugation and incubated at 95°C for 10 minutes. Next, 10μg of lysate was added to 200μL urea buffer (8M urea, 0.1M Tris/HCl pH 8.5) and loaded into 30 KD Vivacon 500 filter units (Saratorius), and centrifuged at 13000g for 15 minutes, and then the concentrated protein was washed three times with urea buffer. The concentrate was alkylated with 50 mM iodoacetamide in urea buffer and incubated in the dark at room temperature for 20 minutes, followed by centrifugation for 15 minutes. The concentrate was washed twice with urea buffer followed by two washes with 50 mM ammonium bicarbonate. Next, 10μg of protein was subjected to trypsin digestion, added at a 1:40 enzyme ratio and incubated for 18 hours at 37°C. The peptides were then collected by centrifugation at 13000g for 15 minutes. The filters were washed with 50 mM ammonium bicarbonate and the wash was also collected by centrifuging at 13000g for 15 minutes. The collected peptides were acidified to 1% formic acid and placed into mass spectrometry vials for analysis.

### Mass Spectrometry and Data Analysis

Tryptic peptides were analyzed as previously published^16,21,43^ by nanoflow LC-MS/MS on a Thermo Orbitrap Velos Pro interfaced with a Thermo EASY-nLC 1000 reversed-phase column (75µm inner diameter 15cm, Reprosil C-18 AQUA 3µm particle size; New Objective) and a flow rate of 400 nl/minute. For peptide separation, a multi-step gradient was utilized from 98% buffer A (0.1% formic acid, 5% DMSO) and 2% buffer B (0.1% formic acid, 5% DMSO in acetonitrile) to 10% buffer A and 90% buffer B over 180 minutes. The spectra were acquired using N^th^ order double-play, data-dependent acquisition mode for the top 20 most abundant ions in the parent spectra for fragmentation. MS1 scans were acquired in Orbitrap mass analyzer at a resolution of 30000. MS1 ions were fragmented by CID fragmentation with an activation time of 30 ms and normalized collision energy 30. Dynamic Exclusion was enabled to avoid multiple fragmentations of parent ions.

The raw files generated were searched against the Uniprot mouse database using Proteome Discoverer 2.5 interfaced with Sequest HT or MaxQuant v1.6.7.0 interfaced with the Andromeda search engine. Subsequent analysis was performed in Proteome Discoverer 2.5 or Perseus v1.6.5.0. The peptides were searched for the static modification of carbamidomethylation on cysteine and the variable modifications of oxidation (M). Proteins with a log_2_fold change >1.5 and an adjusted *p*-value <0.05 were considered as differentially regulated proteins. Gene ontology (GO) analysis for differentially regulated proteins was conducted using the Database for Annotation, Visualization, and Integrated Discovery. When we searched for enrichment of GO terms, Benjamini-Hochberg correction on *p*-value was used to determine GO significance. The mass spectrometry raw files have been uploaded to the PRIDE database via the PRIDE partner repository with the data set identifier PXD.

### Statistical analysis

Unless specifically noted, data were collected from technical triplicates of at least three biological replicates of three independent experiments and presented as the means with standard error of the mean (SEM). Statistical significance was evaluated using the unpaired Student’s *t*-test, one-way or two-way ANOVA for repeated measures, or pairwise comparisons. An asterisk * that indicates *p* <0.05 was considered statistically significant. The data were analyzed using Microsoft Office Excel or GraphPad Prism.

## RESULTS

### SMYD5 expression is dysregulated in the diseased heart

To determine where SMYD5 is expressed, we compiled data from 30 cell and tissue types showing SMYD5’s expression profile from the Human Proteome Map portal^44^ (Figure 1A). We show that SMYD5 is ubiquitously expressed in multiple tissues and hematopoietic cells, with the highest expression in reproductive tissues and immune cells, among others. Next, we examined, for the first time, the intracellular location of SMYD5 and showed that while SMYD5 is expressed in both the cytoplasm and nucleus of HeLa cells, it is more abundant in the cytoplasm of neonatal rat ventricular myocytes (NRVMs). Although upon treatment with the hypertrophic agonist (phenylephrine, PE), SMYD5 translocated to the nucleus of NRVM cells (Figure 1B). This is consistent with data from other SMYD family members, which are also expressed in the cytosol and nucleus and undergo translocation during stress ^20^.

Next, we utilized two mouse models of cardiac hypertrophy to examine SMYD5 expression in the heart during cardiac stress: pressure overload via transverse aortic constriction (TAC) and isoproterenol (ISO) infusion via mini-osmotic pump implantation. We found that *Smyd5* expression decreases during cardiac hypertrophy by ∼50% and ∼30%, respectively, in these models (Figure 1C). Similarly, in a cell model of hypertrophy, where NRVMs were treated with PE, *Smyd5* transcript expression was also decreased ∼35% (Figure 1D). Furthermore, we examined *SMYD5* expression in cardiac tissue samples harvested from heart failure patients at the time of left ventricular assist device (LVAD) implant (Figure 1E and Figure 1F). After LVAD implant, patients were monitored for up to six months to determine which patients responded positively to unloading therapy or had no change in cardiac function, as measured by ejection fraction (Figure 1E). Analysis of cardiac tissue via RNA-Seq revealed a significant increase in *SMYD5* expression in patients regardless of their capacity for functional recovery via LVAD unloading (Figure 1F). Interestingly, a targeted analysis of SMYD5 expression by RT-qPCR in a smaller subset of patients with different sex-based demographics^36^ showed that patients who responded to LVAD treatment displayed increased *SMYD5* expression (Figure S1). In contrast, patients who did not respond to LVAD unloading maintained reduced *SMYD5* expression (Figure S1). High variability in human samples often reflects underlying biological differences, such as comorbidities, genetic background, sex, etc. therefore additional research will be needed to understand these differences in SMYD5 abundance.

### Generation of inducible, cardiomyocyte-specific Smyd5 (iSmyd5) knockout mouse model

To investigate the role of SMYD5 exclusively in the heart, as SMYD5 is ubiquitously expressed in several tissue types including fibroblasts and immune cells, we engineered a cardiomyocyte-specific, tamoxifen-inducible *Smyd5* knockout (iSmyd5 KO) mouse model (Figure 2A). The iSmyd5 KO mice are homozygous for the floxed *Smyd5* allele and carry a *Cre* recombinase allele with expression driven by the *α-Mhc* promoter (Figure 2B). To induce *Smyd5* deletion and determine how the loss of *Smyd5* affects the adult mouse heart, mice were fed tamoxifen-containing chow (TMX) for five weeks. Experimental iSmyd5 mice carried both the floxed *Smyd5* alleles and the *αMhc*-Mer-Cre-Mer transgene, whereas littermate control mice either carried the floxed *Smyd5* allele without the *Cre* transgene or carried the *Cre* transgene without the floxed *Smyd5* allele. All control mice received the same TMX-containing chow for the same duration, ensuring that neither tamoxifen administration nor the presence of the *Cre* transgene alone accounted for the observed phenotype. We monitored heart function by echocardiography and harvested cardiac tissue at either one, two, or five weeks after the beginning of TMX administration (Figure 2C). We confirmed attenuated expression of *Smyd5* in cardiac tissue at both the transcript (Figure 2D) and protein levels (Figure 2E and Figure 2F). To further confirm that the loss of *Smyd5* was cardiomyocyte-specific, we isolated cardiomyocytes and cardiac fibroblasts from mouse heart tissue and probed for cell-specific markers (*Myh6*, *Col1a1*, and *Vim*) to confirm enrichment (Figure 2G). We also quantified *Smyd5* in these cell populations (Figure 2H) to show that *Smyd5* is only reduced in cardiomyocytes, with no change in fibroblasts. In addition, *Smyd5* expression in skeletal muscle remained unaffected (Figure 2I).

**Figure 2:**
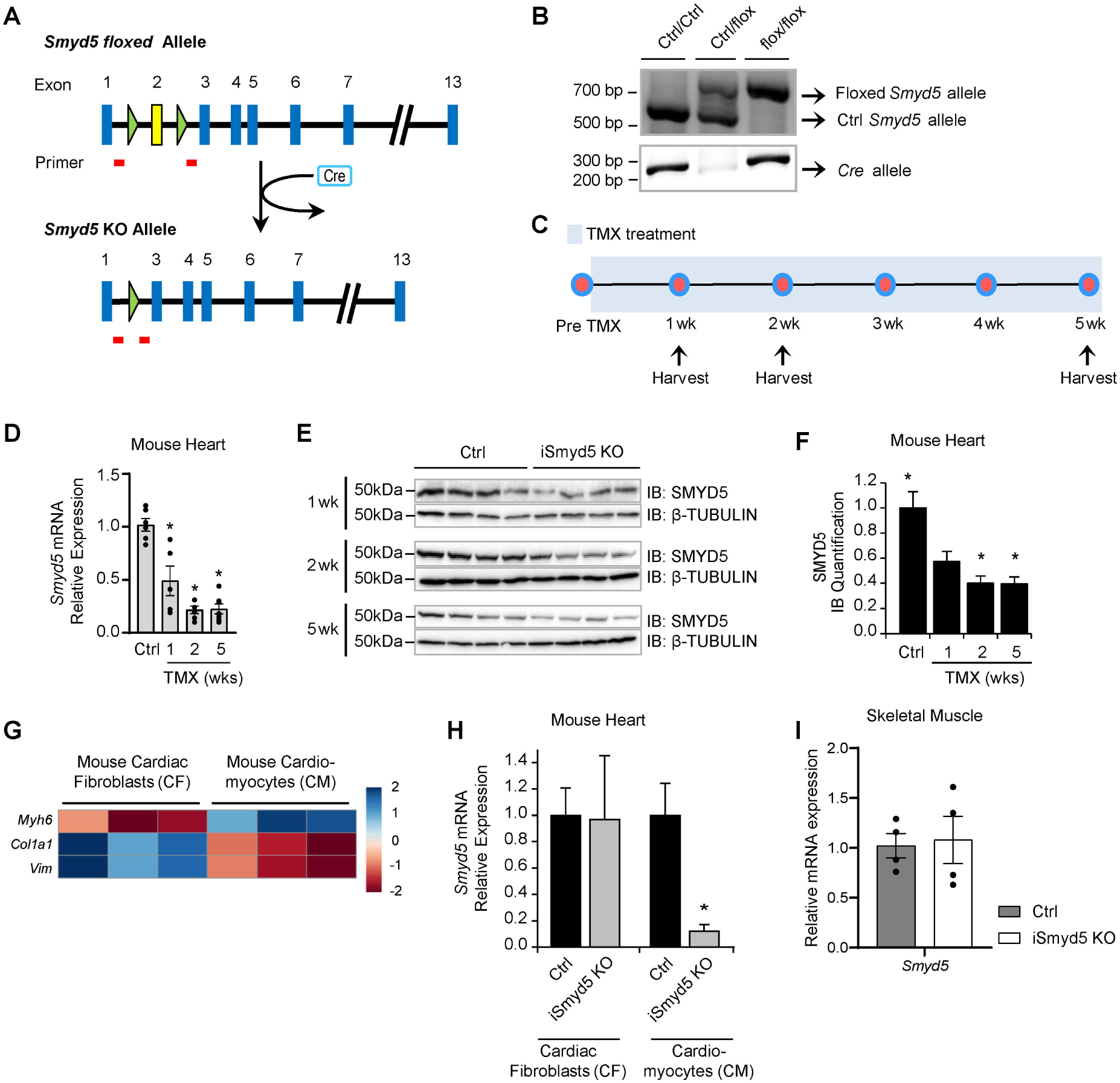
Characterization of cardiac-specific, tamoxifen-inducible *Smyd5* (iSmyd5) knockout mice. **A**, Schematic showing alleles in floxed and iSmyd5 KO mice. Exon 2 is flanked by two *loxP* sites (green triangles). Upon expression of CRE recombinase via the inducible cardiac-specific *Mer-Cre-Mer* allele, exon 2 is excised, resulting in the loss of SMYD5. Red lines indicate primers used for genotyping. **B**, Agarose gel showing the floxed *Smyd5* allele vs. the wild type (Ctrl) *Smyd5* allele, as well as the presence or absence of the *Cre* allele. **C**, Diagram showing the timeline of mice fed tamoxifen (TMX). Cardiac tissue was harvested from mice at either 1, 2, or 5 weeks (wk) after TMX treatment. **D**, Quantification of *Smyd5* expression in cardiac tissue determined via RT-qPCR. *n*=4 (Ctrl), *n*=4 (iSmyd5 KO). A one-way ANOVA was used for statistical analysis. Asterisk * indicates *p*-value <0.05. **E**, Immunoblot analysis of SMYD5 in murine cardiac tissue and **F**, relative expression of protein quantified from immunoblots. *n*=4 (Ctrl), *n*=4 (iSmyd5 KO). A one-way ANOVA was used for statistical analysis. Asterisk * indicates *p*-value <0.001. **G**, Heatmap showing gene expression by RT-qPCR of cell-specific markers for cardiomyocytes and fibroblasts isolated from cardiac tissue. *n*=3 (Ctrl), *n*=3 (iSmyd5 KO). **H**, RT-qPCR analysis of *Smyd5* in mouse cardiac fibroblasts and cardiomyocytes. *n*=3 (Ctrl), *n*=3 (iSmyd5 KO). A two-tailed unpaired Student’s *t*-test was used for statistical analysis. Asterisk * indicates *p*-value <0.01. **I**, Transcript expression of *Smyd5* in skeletal muscle isolated from iSmyd5 KO mice at one week of TMX administration, showing *Smyd5* knockdown is cardiac-specific. *n*=4 (Ctrl), *n*=4 (iSmyd5 KO). A two-tailed unpaired Student’s *t*-test was used for statistical analysis. Data are expressed as the mean ± SEM (**D, F, H, I**).

### Loss of Smyd5 in adult mouse heart induces heart failure

To further evaluate the effects of *Smyd5*’s loss on adult cardiac function, we isolated cardiac tissue from iSmyd5 KO mice at staggered time points to examine physiological and molecular indices. Specifically, we quantified the heart weight-to-body weight ratios, which increased at both two- and five-week time points of TMX treatment, indicating organ-level hypertrophy (Figure 3A). Evaluation of cardiac function by echocardiography showed a significant reduction in the ejection fraction (Figure 3B), fractional shortening (Figure 3C), and stroke volume (Figure 3D) of iSmyd5 KO mice after only one week of TMX treatment, which continued to decline by five weeks. Additional echocardiographic evaluation of left ventricular chamber dimensions revealed progressive increases in both left ventricular end-diastolic diameter (LVEDD, Figure 3E) and left ventricular end-systolic diameter (LVESD, Figure 3F), consistent with progressive left ventricular dilation in iSmyd5 KO mice. Histological evaluation of H&E-stained sections of cardiac tissue from iSmyd5 KO hearts revealed dilated left ventricular chambers at these same time points, two- and five-weeks of TMX treatment (Figure 3G). Trichrome staining of heart tissue displayed an increase in fibrosis as well as increased cell infiltration in the iSmyd5 KO heart tissue (Figure 3H). Quantification of myocytes from cardiac tissue sections stained with wheat germ agglutinin (WGA) showed increased cardiomyocyte size, confirming cellular hypertrophy due to loss of *Smyd5* (Figure 3I and Figure 3J). Next, we examined the transcript abundance of *Nppa*, *Myh6*, *Myh7, Atp2a2,* and the fibrotic marker *Vim*, which revealed significant changes following the loss of *Smyd5 -* changes that are characteristic of cardiac hypertrophy, heart failure, and fibrosis (Figure 3K through 3O).

**Figure 3:**
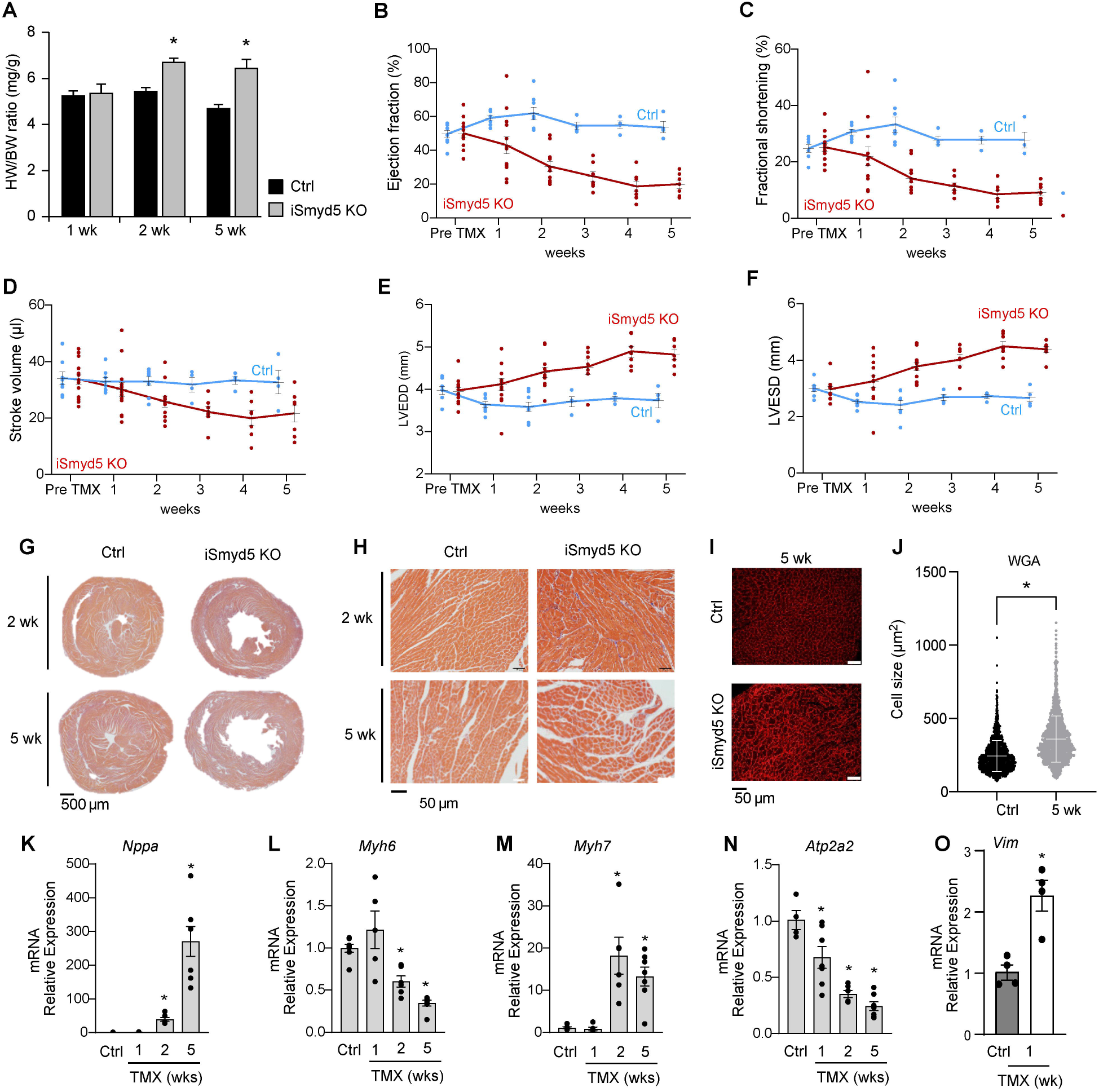
Loss of *Smyd5* in the adult heart induces heart failure. **A**, Loss of *Smyd5* induced cardiac hypertrophy (as measured by heart weight:body weight ratio, HW/BW). *n*=8 (Ctrl), *n*=13 (iSmyd5 KO). A two-way repeated-measures ANOVA was used for statistical analysis. Asterisks * indicate *p*-value <0.005. **B-F**, Cardiac function was measured by electrocardiography and indicated that loss of *Smyd5* induces progressive decline in left ventricular ejection fraction (**B**), fractional shortening (**C**), and stroke volume (**D**), as well as increased left ventricular end-diastolic (**E**), end-systolic (**F**) diameter. iSmyd5 KO mice are indicated in red line, and Ctrl mice are indicated in blue line. Arrows in **B** indicate time points at which cardiac tissue was harvested. *n*=8 (Ctrl), *n*=13 (iSmyd5 KO). A two-way repeated-measures ANOVA was used for statistical analysis. Asterisks * indicate *p*-value <0.005. **G**, Trichrome staining of heart tissue sections obtained from iSmyd5 KO and Ctrl mice, depicting dilated chambers in iSmyd5 KO mice. **H**, Enlarged images from sections depicted in (**G**) show fibrosis after loss of *Smyd5*. *n*=4 (Ctrl), *n*=4 (iSmyd5 KO) at each time point. Representative images are shown. **I**, WGA staining and **J**, quantification of cell size in tissue sections from iSmyd5 KO and Ctrl hearts. *n*=4 (Ctrl), *n*=4 (iSmyd5 KO). Fifty cells were quantified in four different field of vies in each biological replicate at week 5. A two-tailed unpaired Student’s *t*-test was used for statistical analysis. Asterisk * indicates *p*-value <0.01. **K-O**, Quantitation of cardiac genes and the fibrotic marker vimentin determined via RT-qPCR. *n*=3. A one-way repeated-measures ANOVA was used for statistical analysis. Asterisk * indicates *p*-value <0.05. Data are expressed as the mean ± SEM (**A-F, J-O**).

We also quantified the transcript (Figure S2A) and protein (Figure S2B through S2G) expression of other SMYD family members, via RT-qPCR and immunoblotting, respectively, and found that only *Smyd2* was upregulated at an early time point concomitant with *Smyd1* being downregulated.

### SMYD5 directly regulates cytokine production in cardiomyocytes

Histopathological scoring of cardiac tissue sections from iSmyd5 KO mice was performed by a board-certified veterinary pathologist (ACVP) using a standardized semi-quantitative ranking scale, as described in the *Methods*. Visual inspection revealed evidence of inflammatory remodeling, which was subsequently quantified using established indices. Detailed histological analysis of cardiac tissue from iSmyd5 KO mice at two- and five-week time points demonstrated the presence of mononuclear cell infiltrates, interstitial fibrosis, and increased heterogeneity in cardiomyocyte cross-sectional areas (Figure 4A through 4D). Inflammatory infiltrates consisted primarily of lymphocytes and macrophages, with rare polymorphonuclear neutrophils, localized within interstitial regions and perivascular areas. These infiltrative lesions were often accompanied by increased collagen deposition, visualized as blue staining in Masson’s trichrome sections, and were predominantly observed in the left ventricular myocardium.

**Figure 4:**
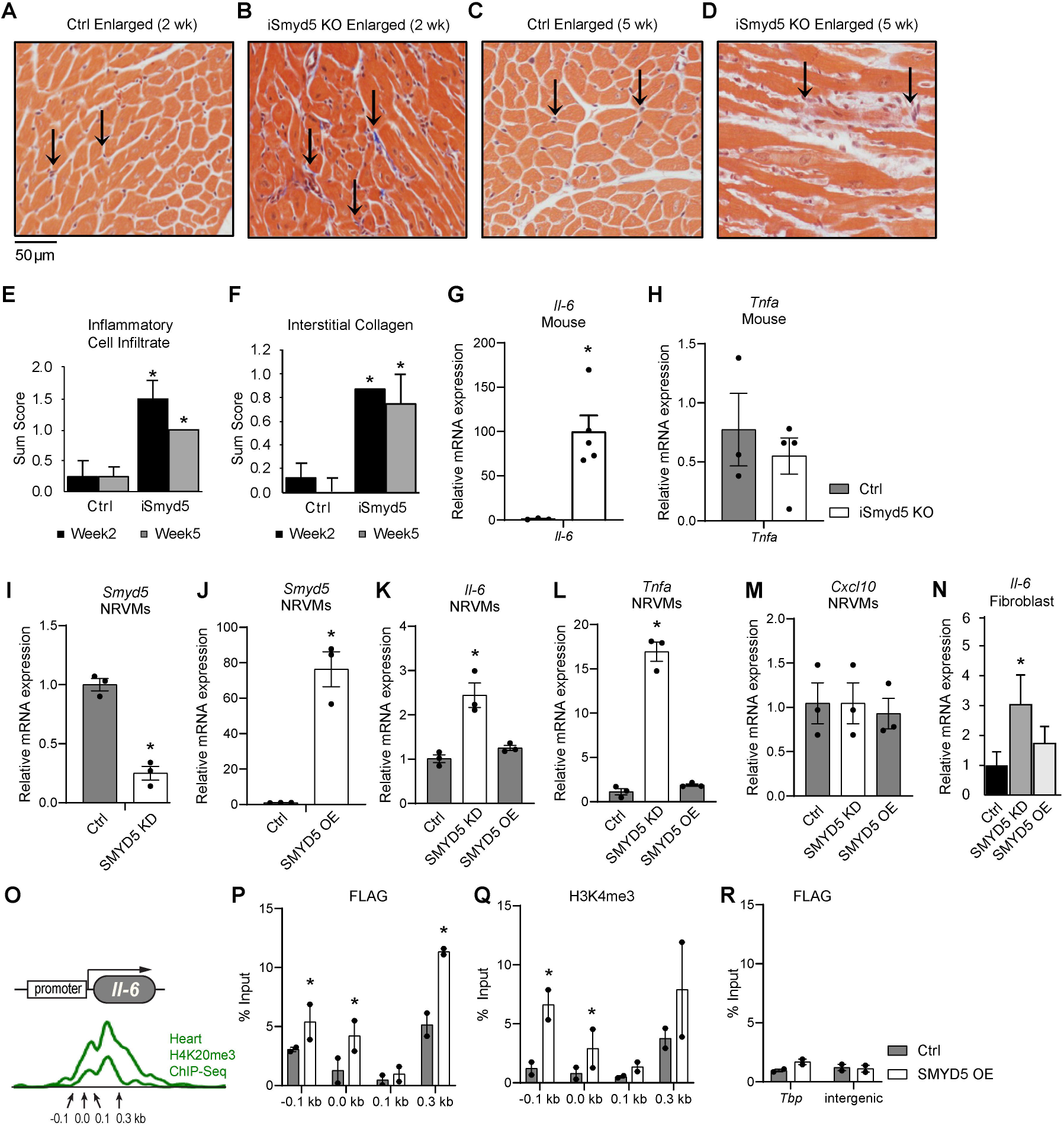
Loss of *Smyd5* leads to increased cytokine production and inflammatory cell infiltration via direct regulation of *Il-6*. **A-D**, High-resolution histology images of Ctrl and iSmyd5 KO hearts stained with H&E show inflammatory cell infiltration (arrows). Representative images are shown. **E**, Inflammatory cell infiltration and **F**, interstitial collagen were quantified using Masson’s trichrome-stained tissue sections; scores were summed for each animal. *n*=4 (Ctrl), *n*=4 (iSmyd5 KO). **G-H**, Quantitation of (**G**) *Il-6* and (**H**) *Tnfα* transcript expression at one-week post-knockout in iSmyd5 KO mice. *n*=4 (Ctrl), *n*=4 (iSmyd5 KO). **I-J**, *Smyd5* transcript expression was quantified in NRVMs after (**I**) *Smyd5* knockdown (KD) via siRNA and (**J**) SMYD5 overexpression (OE) via adenovirus. *n*=3. **K-L**, Quantitation of (**K**) *Il-6* and (**L**) *Tnfα* in NRVMs after *Smyd5* KD, or SMYD5 OE. *n*=3. **M**, Quantitation of *Cxcl10* in NRVMs and (**N**) *Il-6* in cardiac fibroblasts after *Smyd5* KD or SMYD5 OE. A two-tailed unpaired Student’s *t*-test was used in **E-N** for statistical analysis. Asterisks * indicate *p*-value <0.05. **O**, Previous ChIP-Seq studies provided enrichment data for histone H3K4me3 at the *Il-6* promoter, which was used to design primers for qPCR. ChIP-qPCR for adenovirus-mediated SMYD5 OE (using FLAG antibody) shows (**P**) enrichment at the promoter region of *Il-6* and (**Q**) increased trimethylation of histone H4K20. **R**, The *Tbp* promoter and an intergenic region were used as negative controls to show that SMYD5 is not enriched at these regions. *n*=2 (confirmed in multiple independent experiments). A two-way repeated-measures ANOVA was used for statistical analysis. Asterisk * indicates *p*-value <0.05. Data are expressed as the mean ± SEM (**E-N, P-R**).

Severity of infiltration ranged from minimal (1–10% affected area) to mild (10–20%), while collagen deposition was consistently scored as minimal across all groups. Quantitative scoring revealed that iSmyd5 KO mice exhibited 7.5-fold and 5-fold increases in inflammatory infiltrate scores, and 9-fold and 7-fold increases in collagen deposition, compared to wild-type (WT) controls at the two- and five-week time points, respectively (Figure 4E and Figure 4F).

Indeed, previous work has shown that SMYD5 regulates inflammatory signaling in cultured macrophages and synoviocytes, specifically *Il-6* and *Tnfα*^25,28^, therefore, we examined these cytokines in more detail. Interestingly, in cardiac tissue, *Il-6* expression was selectively increased 103-fold at the one-week time point in iSmyd5 KO (Figure 4G), with no change in *Tnfα* expression (Figure 4H). To further evaluate SMYD5’s ability to regulate *Il-6* in cardiomyocytes, we utilized isolated cultured neonatal rat ventricular myocytes (NRVMs) and primary fibroblasts using gain- and loss-of-function analyses (Figure 4I and Figure 4J). In NRVMs, loss of *Smyd5* increased *Il-6* (by ∼2.5-fold) and *Tnfα* (by ∼17-fold) expression, while overexpression of SMYD5 maintained abundance near physiological levels (Figure 4K and Figure 4L), similarly to the *Cxcl10* levels evaluated in these conditions (Figure 4M). In primary fibroblasts, *Il-6* expression also increased in response to *Smyd5* knockdown (Figure 4N), consistent with previous reports in fibroblast cells^28^.

To determine if SMYD5 directly regulates *Il-6* expression in the cardiomyocyte via histone modification of the promoter, we performed chromatin immunoprecipitation of FLAG-tagged SMYD5 in isolated cardiomyocytes. Immunoprecipitated DNA was analyzed by ChIP-qPCR in the promoter region for *Il-6,* as indicated in Figure 4O, with primers developed at sites indicated by arrows (Figure 4O). We detected increased binding of SMYD5 at the promoter (Figure 4P) concomitant with increased enrichment of histone H4K20me3 at these same sites in the *Il-6* promoter (Figure 4Q) and no enrichment in the negative control regions (Figure 4R). Collectively, this data shows for the first time that SMYD5 is an epigenetic regulator of intrinsic cardiomyocyte regulation of inflammatory signaling.

### SMYD5 regulates histone post-translational modifications

Previous studies have identified SMYD5 as a histone methyltransferase capable of catalyzing methylation of histone H4 at lysine K20 (H4K20), both in vitro and in cultured macrophages or embryonic stem cells^24,25^. Among the histone modifications reported to be regulated by SMYD5, histone H4K20 methylation is the most consistently observed across multiple experimental systems and is therefore considered its best-established substrate^23–25^. In contrast, methylation of histone H4 at lysine 36 (H3K36) and lysine 37 (H3K37) has only been reported under specific experimental conditions and has not been consistently reproduced across studies, suggesting that these modifications may be context-dependent^29,30^. Based on this evidence, we focused our analyses on histone H4K20 methylation, which is the primary epigenetic mark regulated by SMYD5. Our targeted analysis of the *Il-6* promoter confirmed reduced histone H4K20 methylation following *Smyd5* deletion in cardiomyocytes. Therefore, we were next interested in determining whether loss of *Smyd5* similarly altered global histone methylation in cardiac tissue. In addition to histone H4K20 trimethylation, we examined histone H3K4me3 and H3K9me3, two well-established histone modifications associated with transcriptionally active euchromatin and repressive heterochromatin, respectively^45–47^. Our immunoblot analyses revealed a significant reduction in global histone H4K20 trimethylation in iSmyd5 KO mouse hearts after one or two weeks of TMX treatment (Figure 5A and Figure 5B). We also observed an increase in histone H3K4me3 after two weeks of TMX treatment, consistent with enhanced transcriptional activation, whereas histone H3K9me3 levels were significantly increased at the five-week time point, suggesting the establishment of a compensatory repressive chromatin state (Figure 5A and Figure 5B).

**Figure 5:**
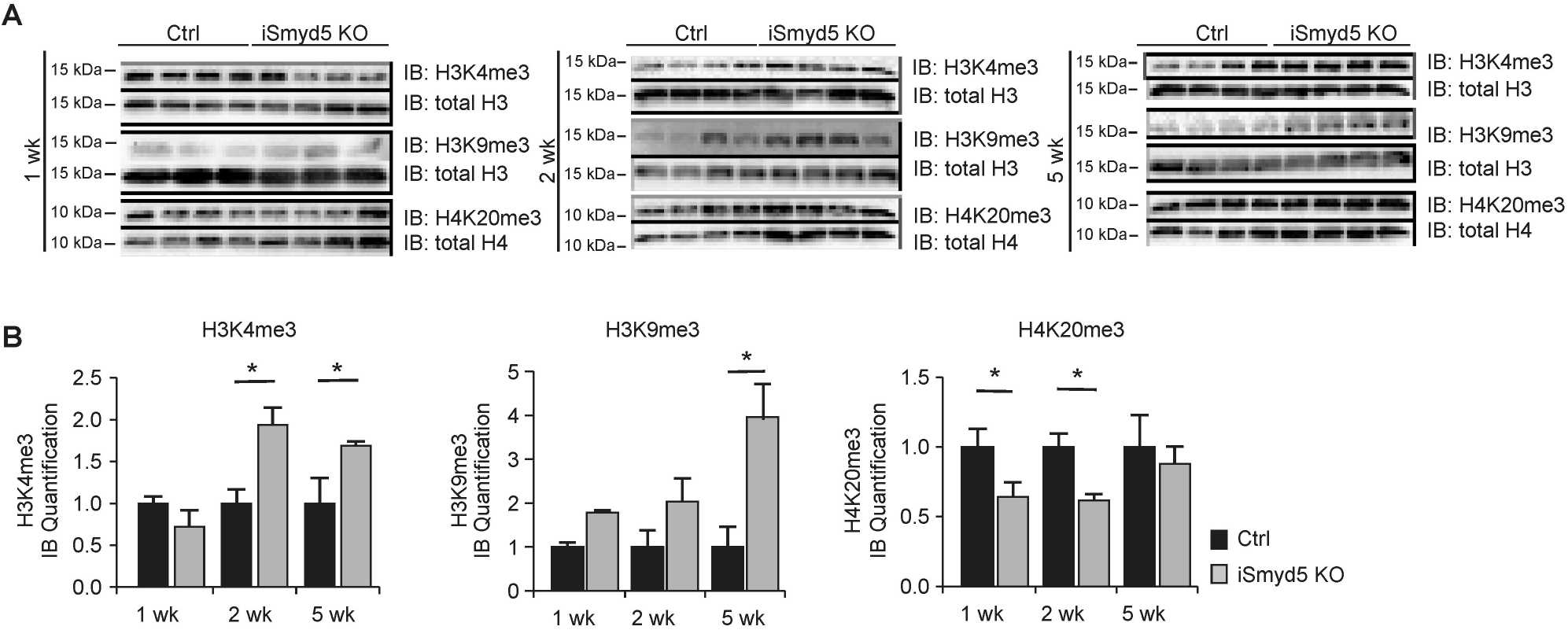
Loss of *Smyd5* in the adult heart affects global histone PTMs. **A**, Trimethylation of histone H4 lysine 20 and histone H3 lysine 4 and lysine 9 in cardiac tissue via immunoblot. **B**, Quantitation of post-translational modifications from immunoblots in panel **A**. *n*=4. A two-tailed unpaired Student’s *t*-test was used for statistical analysis. Asterisk * indicates *p*-value <0.05. Data are expressed as the mean ± SEM (**B**).

### Proteomics analysis of iSmyd5 mouse cardiac tissue

To investigate how loss of *Smyd5* alters protein expression profiles in the heart, we performed label-free quantitative (LFQ) LC-MS/MS proteomics on ventricular tissue from iSmyd5 KO mice and their littermate controls following one, two, and five weeks of TMX treatment. Proteins were identified and quantified using stringent criteria, requiring at least two unique peptides per protein, a high protein-level false discovery rate (FDR) confidence score, and the removal of known contaminants. In total, 1,667 proteins met these inclusion thresholds and were used for downstream analysis. A principal component analysis (PCA) plot generated from all quantified proteins (Figure S3A) revealed clear separation between KO and control tissue at all time points, confirming both dataset reproducibility and distinct proteomic signatures induced by *Smyd5* loss. The expression patterns of all 1,667 quantified proteins in iSmyd5 KO hearts at one, two, and five weeks are displayed in a heatmap (Figure S3B) where protein intensities were log₂-transformed, normalized, and subjected to hierarchical clustering. Protein abundances for each biological replicate were averaged by time point, resulting in a single representative column per KO time point. The heatmap reveals progressive remodeling of the proteome, with distinct clusters of proteins showing either early induction or suppression at one week, followed by broader changes at two and five weeks. These patterns indicate temporal reprogramming of the cardiac proteome after loss of *Smyd5*. For differential expression analysis, protein intensities were background-corrected, and missing values for low-abundance proteins were imputed by low-abundance resampling. Protein ratios were calculated using a pairwise ratio-based approach comparing KO versus control samples at each time point. Hypothesis testing was performed using a background-based two-sided *t*-test, and proteins were considered differentially expressed if they passed both the fold change (log₂FC >1.5) and *p*-value (*p* <0.05) thresholds after filtering. This statistical approach allowed us to robustly identify proteins whose changes were both biologically meaningful and statistically significant. Using these criteria, we identified 85, 142, and 118 differentially expressed proteins at one, two, and five weeks, respectively (Figure S3C). The Venn diagram highlights the overlap between differentially expressed proteins at different time points, showing both shared and time-specific alterations.

We next focused on the one-week time point, representing an early stage where *Smyd5* loss has occurred but precedes heart failure, thus capturing formative changes most directly attributable to *Smyd5* deletion. Expression data at one week for all 1,667 proteins is visualized in a volcano plot (Figure S3D), which displays the statistical significance (–log₁₀ *p*-value) versus magnitude of change (log₂ fold change). Proteins meeting the expression thresholds are presented as dark circles, with Growth Factor Receptor-Bound Protein 2 (GRB2) and Signal Transducer and Activator of Transcription 3 (STAT3) pseudo colored red as they are established downstream targets of *Il-6*.To further define the pathways affected by *Smyd5* deficiency, we performed gene ontology (GO) analysis on the differentially expressed proteins at one week (Figure S3E), which included enrichment for proteins involved in the regulation of gene expression, cellular response to interleukin, and response to cytokine, consistent with activation of inflammatory signaling pathways. To examine the abundance of GRB2 and STAT3, box-and-whisker plots were generated (Figure S3F through S3H) showing a ∼24-fold (log₂FC ≈ 4.5) and ∼3-fold (log₂FC ≈ 1.6) increase, respectively. These box-and-whisker plots show the distribution of LFQ intensities across biological replicates, whereas the line plot (Figure S3I) illustrates the dynamic time course of GRB2 expression.

Among the notable proteins identified, STAT3 plays a central role in cytokine signaling and inflammation^48,49^ as a known downstream target of IL-6^50^, and its early upregulation in iSmyd5 KO hearts (Figure S3F) supports activation of IL-6–mediated pathways. Similarly, GRB2, an adapter protein that links receptor signaling to multiple downstream cascades, was one of the most upregulated proteins at one week (Figure S3G through S3I). GRB2 regulates diverse biological processes, including innate and adaptive immunity^51,52^, and as a downstream target of IL-6^53^, its expression has been associated with detrimental effects on cardiomyocytes and diastolic dysfunction^51^. Together, these findings suggest that loss of *Smyd5* rapidly activates IL-6–dependent inflammatory signaling, with STAT3 and GRB2 emerging as central mediators of the early cardiac response to *Smyd5* deletion.

### SMYD5 overexpression attenuates cellular hypertrophy in vivo

Because the loss of *Smyd5* induces inflammation but also cardiac pathological remodeling that progresses to heart failure in adult mice, we next sought to determine the effects of *Smyd5* overexpression in cardiomyocytes. Utilizing cultured cells, we confirmed the efficacy of an adenovirus expressing FLAG-tagged SMYD5 (SMYD5-OE) or Null (control) virus in H9c2 cells (Figure 6A). Similar data were also generated in NRVM cells. Adenovirus-mediated overexpression of SMYD5 did not affect the basal cell phenotype; however, overexpression of SMYD5 increased global levels of histone H4K20 trimethylation (Figure 6A), and attenuated PE-induced hypertrophic growth (Figure 6B and Figure 6C).

**Figure 6:**
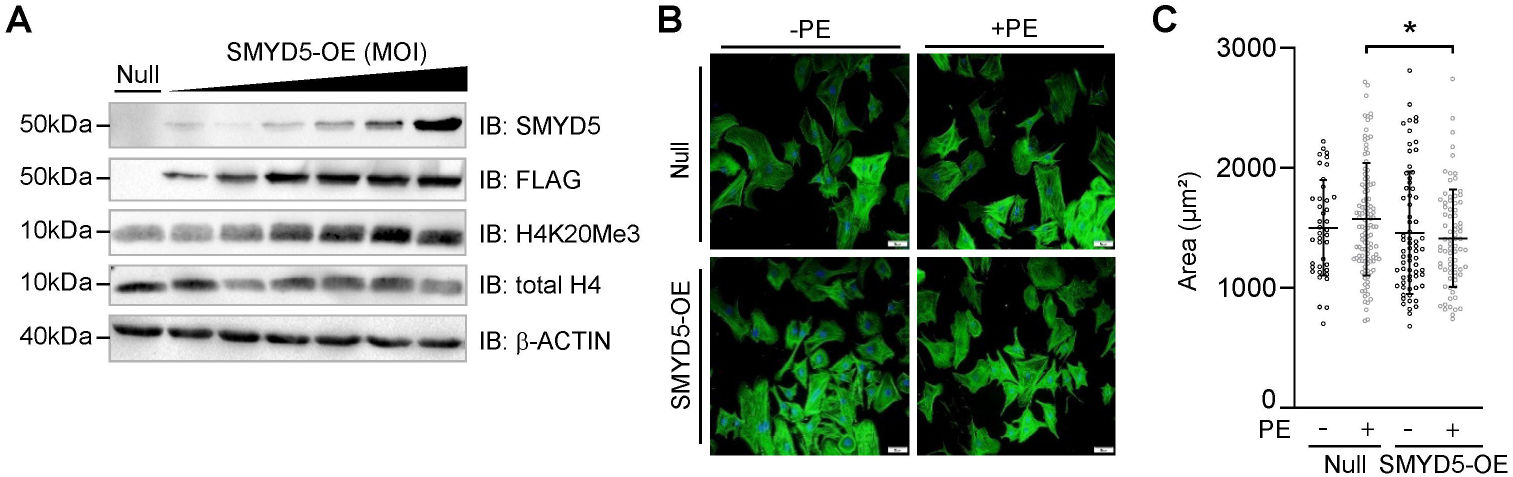
Overexpression of SMYD5 inhibits hypertrophic growth in cardiomyocytes. **A**, Adenoviral transduction of H9c2 rat cardiomyocytes led to robust expression of FLAG-tagged SMYD5, as confirmed by immunoblotting. MOI, multiplicity of infection. Null, empty virus. Increasing SMYD5 leads to an increase in histone H4K20me3. **B**, Neonatal rat cardiomyocytes were transduced with Null or SMYD5 adenovirus in the presence and absence of PE for 48 hours, stained with phalloidin and analyzed using fluorescent microscopy. **C**, Cardiomyocyte size was quantified and showed that overexpression of SMYD5 inhibits PE-induced hypertrophic growth. *n*=40 (Null), *n*=114 (Null+PE), *n*=69 (SMYD5-OE), *n*=78 (SMYD5-OE+PE), quantified from three independent experiments. A two-way ANOVA with a post-hoc Tukey test for multiple comparisons was used for statistical analysis. Asterisk * indicates *p*-value <0.05. Data are expressed as the mean ± SEM (**C**).

## DISCUSSION

In this study, we generated the first mouse model to evaluate the role of SMYD5 in the heart and demonstrate that SMYD5 is a novel epigenetic regulator of *Il-6* expression and intrinsic inflammatory signaling in the cardiomyocyte. By analyzing the loss-of-function model, we showed that cardiomyocyte-specific ablation of *Smyd5* is sufficient to induce pathological cardiac remodeling characterized by robust inflammatory signaling as well as hypertrophic growth, which progresses to fulminant heart failure. This phenotype, which exhibited features akin to myocarditis or cytokine storm-like syndromes, was accompanied by mononuclear cell infiltration into cardiac tissue and a significant increase in *Il-6* expression, along with its downstream targets. We propose that during normal cardiac function, SMYD5 represses *Il-6* in cardiomyocytes via trimethylation of histone H4K20. However, cardiomyocyte-specific loss of *Smyd5* leads to a massive increase in *Il-6* expression and activation of inflammatory signaling pathways, hypertrophic growth, increased fibrosis, ultimately progressing to cardiac dysfunction and heart failure (Figure 7). Under normal, basal conditions, healthy cardiomyocytes produce little to no detectable *Il-6* mRNA expression and relatively low levels of IL-6 protein, whereas IL-6 expression is strongly upregulated in response to cellular stress or injury^54,55^. While IL-6 plays a protective role in the acute response to injury by activating immune cells and initiating healing processes, persistent elevation of IL-6 leads to chronic inflammation and fibrosis^56^. This dual role of IL-6 in response to injury is particularly apparent in cardiac tissue. Specifically, previously published studies utilizing animal models of cardiac disease showed that short-term IL-6 signaling protects cardiac tissue in response to acute damage^57–59^. In contrast, long-term IL-6 signaling is associated with maladaptive hypertrophy and decreased contractile function, which progresses to cardiovascular disease^60,61^. Indeed, transgenic mice overexpressing IL-6/IL-6R develop ventricular cardiac hypertrophy^62^. Consistent with these models, elevated *IL-6* has been reported as a predictor of new-onset heart failure and is upregulated in human heart failure cohorts, as measured in serum samples collected from a prospective general population-based cohort study, PREVEND^63^.

**Figure 7:**
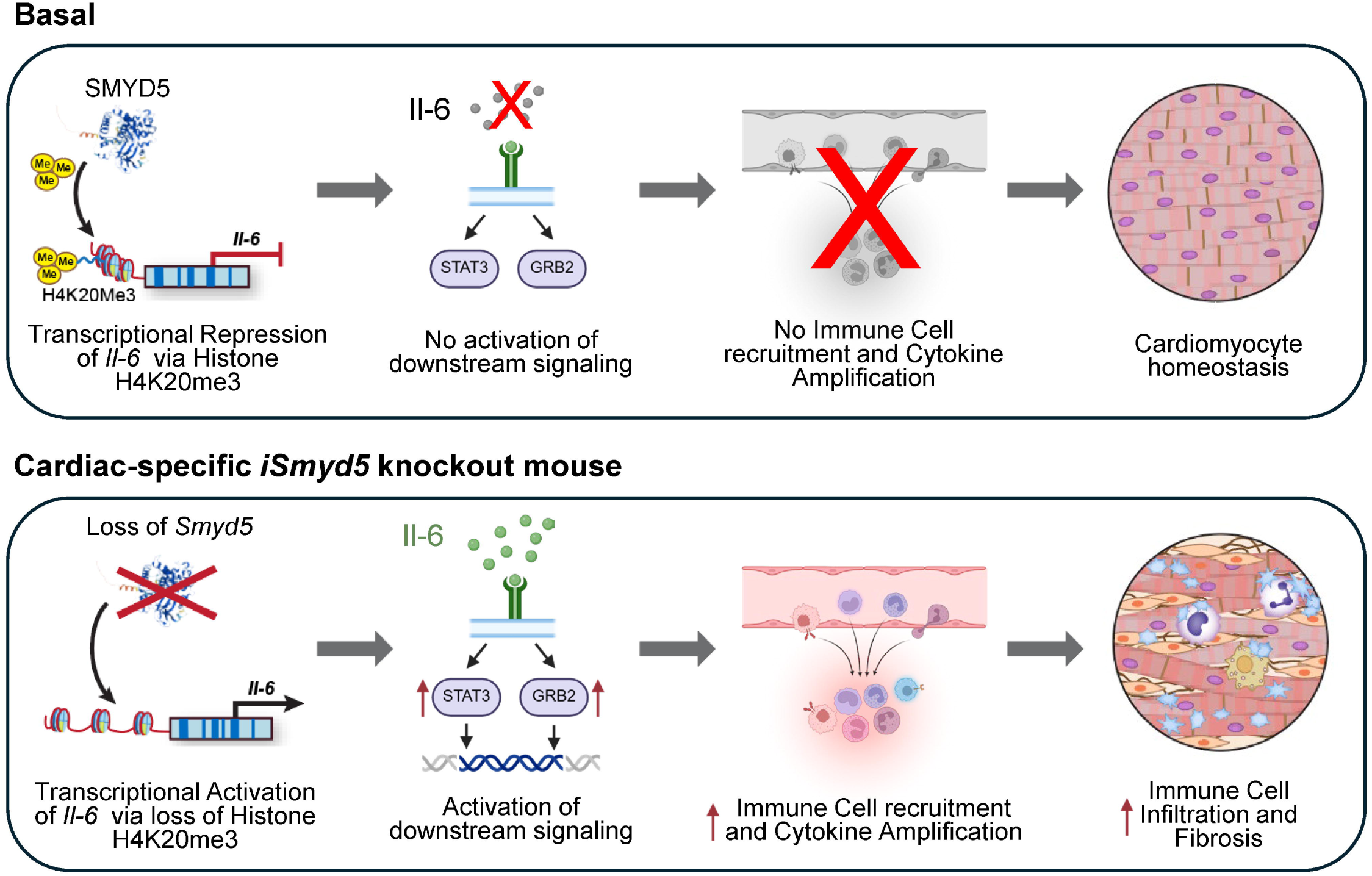
SMYD5 regulates cardiac inflammatory response in cardiomyocyte. Upper panel: Under basal conditions, SMYD5 inhibits intrinsic cardiomyocyte expression of inflammatory cytokines (i.e. *Il-6*) via histone H4K20 trimethylation of the promoter resulting in maintained cardiomyocyte homeostasis. **Lower panel:** Cardiac-specific loss of *Smyd5* leads to transcriptional activation of *Il-6* and activation of downstream signaling leading to immune cell recruitment, cytokine amplification, and robust induction of immune cell infiltration and fibrosis in the heart.

It is interesting and important to note that this robust inflammatory phenotype is unique to SMYD5 and not observed in knockout models of any other SMYD family members^20,64,65^. In addition, our findings reveal that *Smyd5* knockout causes an unprecedented 100-fold increase in *Il-6* expression, far exceeding the levels examined in transgenic mouse models or observed in human heart failure serum samples (5-10 fold increase)^66,67^. These elevated *Il-6* levels are likely due to amplification of the initial cardiomyocyte signal by invading immune cells or fibroblasts; however, such robust IL-6 levels have only been reported in cases of severe myocarditis or cytokine storm syndromes. Consistent with this inflammatory phenotype, our proteomics analysis identified significant upregulation of proteins involved in IL-6-associated signaling, including GRB2 and STAT3, both of which play central roles in cytokine signaling and have been implicated in cardiac dysfunction^68^. Notably, STAT3 is constitutively expressed in healthy cardiomyocytes under normal physiological conditions, where it contributes to cellular homeostasis and cardiac protection^69,70^. GRB2 and STAT3 are also established components of intracellular signaling pathways, with GRB2 serving as a broadly acting adaptor protein that integrates signals from multiple receptors^51,52^. Thus, the increased abundance of these proteins in *Smyd5*-deficient hearts may reflect enhanced inflammatory signaling rather than their de novo expression. Although inflammatory signaling is markedly enhanced in *Smyd5*-deficient hearts and likely contributes to the observed cardiac dysfunction, we cannot exclude the possibility that the phenotype also reflects cardiomyocyte-intrinsic dysregulation independent of, or synergistic with, cytokine signaling. Given the epigenetic regulatory function of SMYD5, its loss may disrupt additional transcriptional programs that directly impair myocardial function while also amplifying inflammatory responses. Thus, although our data support a strong association between inflammation and heart failure, they do not establish a direct causal relationship, raising the possibility that inflammation, cardiac dysfunction, and other SMYD5-dependent mechanisms represent interconnected downstream consequences of *Smyd5* deficiency (Figure 8). Future studies employing genetic or pharmacological inhibition of cardiomyocyte IL-6 signaling in iSmyd5 KO mice will be essential to establish whether the suppression of this inflammatory pathway is sufficient to attenuate cardiac remodeling and preserve cardiac function, thereby distinguishing whether inflammatory activation is a primary driver of heart failure or a parallel consequence of *Smyd5* deficiency.

**Figure 8.**
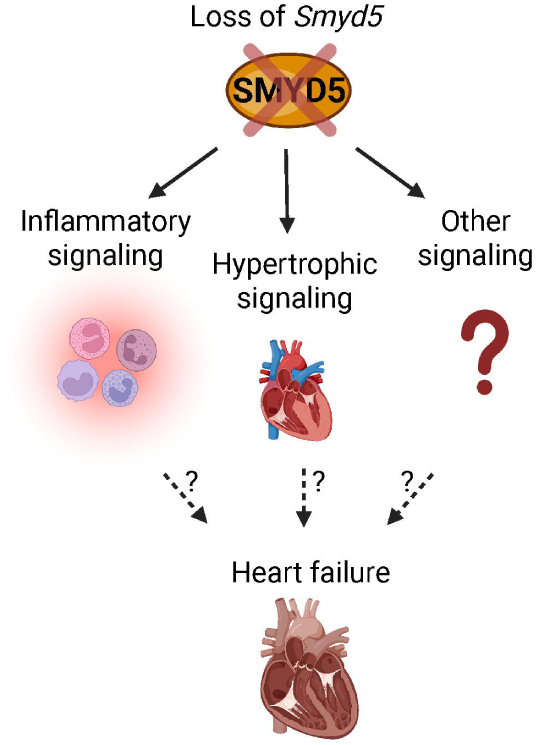
Proposed model of the role of SMYD5 in cardiac homeostasis. Our data demonstrate that loss of *Smyd5* leads to robust inflammation and heart failure in mice. However, the mechanistic relationship between these phenotypes remains unclear. Inflammation may directly drive the development of heart failure, or inflammation and cardiac dysfunction may represent parallel downstream consequences of *Smyd5* deficiency. Alternatively, unidentified mechanisms may contribute to the development of heart failure following *Smyd5* loss.

Collectively, this study identifies SMYD5 as the first epigenetic regulator of intrinsic cardiomyocyte inflammatory signaling and establishes the molecular basis for direct regulation of *Il-6* via SMYD5’s methyltransferase activity. While this work reports the first direct gene target of SMYD5 in the cardiomyocyte, the full profile of genes regulated by SMYD5 in the cardiac genome is unknown. It is also likely that the anti-hypertrophic effects of SMYD5 are not mediated exclusively through *ll-6* repression. Given its broad role in chromatin regulation, SMYD5 may control additional transcriptional programs involved in cardiomyocyte growth and stress adaptation. Defining the direct genomic targets of SMYD5 and functionally interrogating these candidate downstream effectors will be an important focus of future studies. Only one study has previously examined global SMYD5 binding regions via ChIP-Seq in macrophages, which demonstrated that SMYD5 bound and trimethylated histone H4K20 at long interspersed elements (LINE) and long-terminal repeat (LTR) regions in these cells^23^. These LINE and TLR regions are typically heterochromatic and constitutively silenced, although decreased repression of these transposable elements can induce pathogen-free inflammation in cells^71^. To date, no studies have reported genome-wide mapping of SMYD5 binding or histone methyltransferase activity (using ChIP-Seq methods) in cardiomyocytes. Therefore, future work using ChIP-Seq approaches will be necessary to define the full repertoire of genes and chromatin regions regulated by SMYD5.

Together, our data establish SMYD5 as a critical epigenetic regulator that restrains inflammatory signaling in the normal heart and demonstrate how its loss precipitates a cascade of cytokine-mediated responses. These findings further raise the intriguing possibility that individuals with heightened susceptibility to myocarditis or cytokine storm–like syndromes may harbor genetic variants in SMYD5 that impair its regulatory function, warranting investigation into whether alterations in SMYD5 contribute to inter-individual differences in inflammatory vulnerability. In addition, these findings suggest that strategies aimed at preserving or restoring SMYD5 expression may represent a therapeutic approach to attenuate aberrant cytokine signaling.

## DATA AVAILABILITY

The mass spectrometry raw files have been uploaded to PRIDE database via the PRIDE partner repository with the dataset identifier PXD.

## ACKNOWLEDGEMENTS

We thank Huntsman Cancer Institute Biorepository and Molecular Pathology Research Histology Core, Chris Hunter, for his assistance in isolating the NRVMs, and Diana Lim for assisting with the graphical design of the final figures.

## AUTHORS CONTRIBUTIONS

M.R.M. R.B. S.H. and S.F. conception and design of research; M.R.M., R.B., S.H, C.C.L., L.W., E.H., M.W.S., A.A., J.G.L., L.F., and K.D. performed experiments; J.R.V., K.S., C.K. and S.G.D. acquisition and analysis of human clinical data; M.R.M., R.B., S.H., M.W.S., and S.F. interpreted results of experiments; M.R.M, R.B., S.H., M.W.S., and S.F. prepared figures; M.R.M., R.B., S.H., and S.F. drafted the manuscript; M.R.M., R.B., S.H., M.W.S. and S.F. edited and revised the manuscript; M.R.M., R.B., S.H., C.C.L., L.W., E.H., M.W.S., A.A., J.G.L., L.F., K.D., S.T.S., and S.F. approved the final version of the manuscript.

## SUPLEMENTAL MATERIAL

**Table S1: Previously published studies on the histone methyltransferase SMYD5.** The data regarding SMYD5 is relatively sparse in the heart, but it has been previously studied in cultured cells, zebrafish, murine arthritis, and cancer, with 17 primary research articles focused on SMYD5 published to date.

**Table S2:**
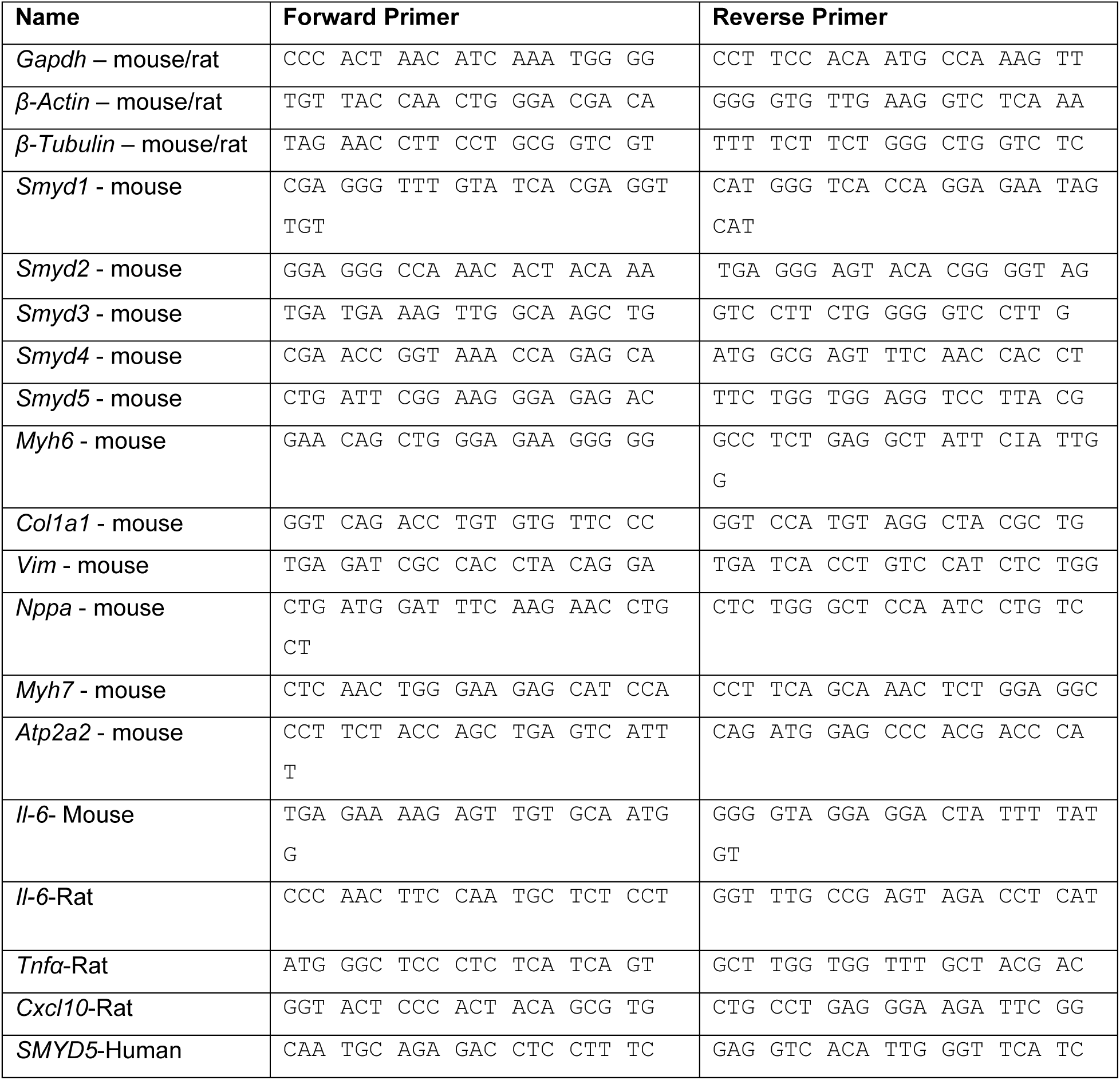
Primers that were used in this study.

**Figure S1: Targeted analysis of *SMYD5* transcript expression in human heart failure patients.** RNA-Seq data showed an increase in *SMYD5* in human heart failure patients in Figure 1. To further evaluate *SMYD5* expression in a targeted manner we designed primers specific to human *SMYD5* and performed RT-qPCR evaluation of its expression in a smaller cohort of human heart failure patients that also underwent LVAD implantation. This cohort, which included higher female representation, showed increased *SMYD5* expression only in patients capable of functional recovery after LVAD unloading (Responders) when compared to non-failing donor (Control) tissue. *n*=4 (Control), *n*=4 (Responders), *n*=3 (Non-responders). A one-way ANOVA was used for the statistical analysis. Asterisk * indicates *p*-value <0.05. Data are expressed as the mean ± SEM.

**Figure S2: Expression of the SMYD family members in the iSmyd5 KO mice. A**, Quantification of transcripts by RT-qPCR in iSmyd5 KO mice. *n*=4. A one-way repeated-measures ANOVA was used for statistical analysis. Asterisk * indicates *p*-value <0.05. **B-G**, Immunoblotting and quantification of SMYD1 (**B,C**), SMYD2 (**D,E**) and SMYD4 (**F,G**) at week 5 in iSmyd5KO mice. *n*=3. A one-way repeated-measures ANOVA was used for statistical analysis. Asterisk * indicates *p* <0.05. Data are expressed as the mean ± SEM (**A, C, E, G**).

**Figure S3: Proteomic analyses of cardiac tissue highlighting changes in inflammatory signaling in iSmyd5 knockout mice. A**, Three-dimensional principal component analysis (PCA) of proteins identified from cardiac tissue harvested from iSmyd5 KO and Ctrl mice at 1, 2, and 5 weeks post-TMX treatment showing distinct clustering of samples, confirming dataset reproducibility and distinct proteomic profiles. **B**, Heatmap generated in Proteome Discoverer 2.5 depicting relative abundance of all 1,667 quantified proteins. Protein abundances were averaged by time point, revealing distinct clustering patterns and progressive remodeling of the proteome. **C**, Venn diagram comparing only differentially expressed proteins (*p*<0.05, log₂ fold change >1.5) at 1, 2, and 5 weeks, illustrating both common and timepoint-specific alterations. **D**, Volcano plot of all 1,667 quantified proteins at 1 week, displaying statistical significance (-log_10_ *p*-value) versus magnitude of change (log_2_ fold change). Significantly upregulated and downregulated proteins (*p*<0.05, log₂ fold change >1.5) are indicated as dark grey circles, GRB2 and STAT3 are pseudo colored red. **E**, Gene ontology (GO) analysis of biological processes enriched in differentially expressed proteins at 1 week post-knockout show strong associations with inflammatory signaling, cytokine response, and gene regulation. Box-and-whisker plot quantifying known downstream targets of Il-6, specifically, **F**, STAT3 abundance at 1 week, and **G–H**, GRB2 abundance at (**G**) 1 week and (**H**) 2 weeks, confirming substantial upregulation (log₂ fold change of ∼4.5 at week 1, and ∼2.2 at week 2). **I**, Line plot showing the temporal trend of GRB2 protein abundance across time points, normalized in Proteome Discoverer 2.5, demonstrating an early peak at 1 week and sustained elevation relative to controls at later stages.

## REFERENCES

1. Hohensinner PJ, Niessner A, Huber K, Weyand CM, Wojta J. Inflammation and cardiac outcome. Curr Opin Infect Dis. 2011;24:259–264. doi: 10.1097/QCO.0b013e328344f50f

2. Roig E, Orus J, Pare C, Azqueta M, Filella X, Perez-Villa F, Heras M, Sanz G. Serum interleukin-6 in congestive heart failure secondary to idiopathic dilated cardiomyopathy. Am J Cardiol. 1998;82:688–690, A688. doi: 10.1016/s0002-9149(98)00388-9

3. Ridker PM, Rifai N, Stampfer MJ, Hennekens CH. Plasma concentration of interleukin-6 and the risk of future myocardial infarction among apparently healthy men. Circulation. 2000;101:1767–1772. doi: 10.1161/01.cir.101.15.1767

4. Edelmann F, Holzendorf V, Wachter R, Nolte K, Schmidt AG, Kraigher-Krainer E, Duvinage A, Unkelbach I, Dungen HD, Tschope C, et al. Galectin-3 in patients with heart failure with preserved ejection fraction: results from the Aldo-DHF trial. Eur J Heart Fail. 2015;17:214–223. doi: 10.1002/ejhf.203

5. Torre-Amione G, Kapadia S, Lee J, Durand JB, Bies RD, Young JB, Mann DL. Tumor necrosis factor-alpha and tumor necrosis factor receptors in the failing human heart. Circulation. 1996;93:704–711. doi: 10.1161/01.cir.93.4.704

6. Vasan RS, Sullivan LM, Roubenoff R, Dinarello CA, Harris T, Benjamin EJ, Sawyer DB, Levy D, Wilson PW, D’Agostino RB, et al. Inflammatory markers and risk of heart failure in elderly subjects without prior myocardial infarction: the Framingham Heart Study. Circulation. 2003;107:1486–1491. doi: 10.1161/01.cir.0000057810.48709.f6

7. D’Elia E, Vaduganathan M, Gori M, Gavazzi A, Butler J, Senni M. Role of biomarkers in cardiac structure phenotyping in heart failure with preserved ejection fraction: critical appraisal and practical use. Eur J Heart Fail. 2015;17:1231–1239. doi: 10.1002/ejhf.430

8. Hinderer S, Schenke-Layland K. Cardiac fibrosis - A short review of causes and therapeutic strategies. Adv Drug Deliv Rev. 2019;146:77–82. doi: 10.1016/j.addr.2019.05.011

9. Fan Z, Guan J. Antifibrotic therapies to control cardiac fibrosis. Biomater Res. 2016;20:13. doi: 10.1186/s40824-016-0060-8

10. Liu T, Song D, Dong J, Zhu P, Liu J, Liu W, Ma X, Zhao L, Ling S. Current Understanding of the Pathophysiology of Myocardial Fibrosis and Its Quantitative Assessment in Heart Failure. Front Physiol. 2017;8:238. doi: 10.3389/fphys.2017.00238

11. Zhao L, Cheng G, Jin R, Afzal MR, Samanta A, Xuan YT, Girgis M, Elias HK, Zhu Y, Davani A, et al. Deletion of Interleukin-6 Attenuates Pressure Overload-Induced Left Ventricular Hypertrophy and Dysfunction. Circ Res. 2016;118:1918– 929. doi: 10.1161/CIRCRESAHA.116.308688

12. Jing R, Long TY, Pan W, Li F, Xie QY. IL-6 knockout ameliorates myocardial remodeling after myocardial infarction by regulating activation of M2 macrophages and fibroblast cells. Eur Rev Med Pharmacol Sci. 2019;23:6283– 6291. doi: 10.26355/eurrev_201907_18450

13. Hang CT, Yang J, Han P, Cheng HL, Shang C, Ashley E, Zhou B, Chang CP. Chromatin regulation by Brg1 underlies heart muscle development and disease. Nature. 2010;466:62–67. doi: 10.1038/nature09130

14. Anand P, Brown JD, Lin CY, Qi J, Zhang R, Artero PC, Alaiti MA, Bullard J, Alazem K, Margulies KB, et al. BET bromodomains mediate transcriptional pause release in heart failure. Cell. 2013;154:569–582. doi: 10.1016/j.cell.2013.07.013

15. Cao DJ, Wang ZV, Battiprolu PK, Jiang N, Morales CR, Kong Y, Rothermel BA, Gillette TG, Hill JA. Histone deacetylase (HDAC) inhibitors attenuate cardiac hypertrophy by suppressing autophagy. Proc Natl Acad Sci U S A. 2011;108:4123–4128. doi: 10.1073/pnas.1015081108

16. Hickenlooper SM, Davis K, Szulik MW, Sheikh H, Miller M, Valdez S, Bia R, Franklin S. Histone H4K20 Trimethylation Is Decreased in Murine Models of Heart Disease. ACS Omega. 2022;7:30710–30719. doi: 10.1021/acsomega.2c00984

17. Hickenlooper S, Brady C, Bia R, Visker JR, Wang L, Valdez S, Gwynn C, Roland MN, Kyriakopoulos CP, Sideris K, et al. Expression profiles of histone H4K20 methylation and its associated enzymes in mouse cardiac disease and human heart failure. Epigenetics. 2025;20:2578553. doi: 10.1080/15592294.2025.2578553

18. Szulik MW, Davis K, Bakhtina A, Azarcon P, Bia R, Horiuchi E, Franklin S. Transcriptional regulation by methyltransferases and their role in the heart: highlighting novel emerging functionality. Am J Physiol Heart Circ Physiol. 2020;319:H847–H865. doi: 10.1152/ajpheart.00382.2020

19. Davis K, Azarcon P, Hickenlooper S, Bia R, Horiuchi E, Szulik MW, Franklin S. The role of demethylases in cardiac development and disease. J Mol Cell Cardiol. 2021;158:89–100. doi: 10.1016/j.yjmcc.2021.05.018

20. Franklin S, Kimball T, Rasmussen TL, Rosa-Garrido M, Chen H, Tran T, Miller MR, Gray R, Jiang S, Ren S, et al. The chromatin-binding protein Smyd1 restricts adult mammalian heart growth. Am J Physiol Heart Circ Physiol. 2016;311:H1234–H1247. doi: 10.1152/ajpheart.00235.2016

21. Warren JS, Tracy CM, Miller MR, Makaju A, Szulik MW, Oka SI, Yuzyuk TN, Cox JE, Kumar A, Lozier BK, et al. Histone methyltransferase Smyd1 regulates mitochondrial energetics in the heart. Proc Natl Acad Sci U S A. 2018;115:E7871–e7880. doi: 10.1073/pnas.1800680115

22. Tracy C, Warren JS, Szulik M, Wang L, Garcia J, Makaju A, Russell K, Miller M, Franklin S. The Smyd Family of Methyltransferases: Role in Cardiac and Skeletal Muscle Physiology and Pathology. Curr Opin Physiol. 2018;1:140–152. doi: 10.1016/j.cophys.2017.10.001

23. Kidder BL, Hu G, Cui K, Zhao K. SMYD5 regulates H4K20me3-marked heterochromatin to safeguard ES cell self-renewal and prevent spurious differentiation. Epigenetics Chromatin. 2017;10:8. doi: 10.1186/s13072-017-0115-7

24. Kidder BL, He R, Wangsa D, Padilla-Nash HM, Bernardo MM, Sheng S, Ried T, Zhao K. SMYD5 Controls Heterochromatin and Chromosome Integrity during Embryonic Stem Cell Differentiation. Cancer Res. 2017;77:6729–6745. doi: 10.1158/0008-5472.CAN-17-0828

25. Stender JD, Pascual G, Liu W, Kaikkonen MU, Do K, Spann NJ, Boutros M, Perrimon N, Rosenfeld MG, Glass CK. Control of proinflammatory gene programs by regulated trimethylation and demethylation of histone H4K20. Mol Cell. 2012;48:28–38. doi: 10.1016/j.molcel.2012.07.020

26. Hou Y, Sun X, Gheinani PT, Guan X, Sharma S, Zhou Y, Jin C, Yang Z, Naren AP, Yin J, et al. Epithelial SMYD5 Exaggerates IBD by Down-regulating Mitochondrial Functions via Post-Translational Control of PGC-1alpha Stability. Cell Mol Gastroenterol Hepatol. 2022;14:375–403. doi: 10.1016/j.jcmgh.2022.05.006

27. Fujii T, Tsunesumi S, Sagara H, Munakata M, Hisaki Y, Sekiya T, Furukawa Y, Sakamoto K, Watanabe S. Smyd5 plays pivotal roles in both primitive and definitive hematopoiesis during zebrafish embryogenesis. Sci Rep. 2016;6:29157. doi: 10.1038/srep29157

28. Xiao C, Su Z, Zhao J, Tan S, He M, Li Y, Liu J, Xu J, Hu Y, Li Z, et al. Novel regulation mechanism of histone methyltransferase SMYD5 in rheumatoid arthritis. Cell Mol Biol Lett. 2025;30:38. doi: 10.1186/s11658-025-00707-9

29. Aljazi MB, Gao Y, Wu Y, He J. SMYD5 is a histone H3-specific methyltransferase mediating mono-methylation of histone H3 lysine 36 and 37. Biochem Biophys Res Commun. 2022;599:142–147. doi: 10.1016/j.bbrc.2022.02.043

30. Zhang Y, Fang Y, Tang Y, Han S, Jia J, Wan X, Chen J, Yuan Y, Zhao B, Fang D. SMYD5 catalyzes histone H3 lysine 36 trimethylation at promoters. Nat Commun. 2022;13:3190. doi: 10.1038/s41467-022-30940-1

31. Park J, Wu J, Szkop KJ, Jeong J, Jovanovic P, Husmann D, Flores NM, Francis JW, Chen YC, Benitez AM, et al. SMYD5 methylation of rpL40 links ribosomal output to gastric cancer. Nature. 2024;632:656–663. doi: 10.1038/s41586-024-07718-0

32. Miao B, Ge L, He C, Wang X, Wu J, Li X, Chen K, Wan J, Xing S, Ren L, et al. SMYD5 is a ribosomal methyltransferase that catalyzes RPL40 lysine methylation to enhance translation output and promote hepatocellular carcinoma. Cell Res. 2024;34:648–660. doi: 10.1038/s41422-024-01013-3

33. Drakos SG, Badolia R, Makaju A, Kyriakopoulos CP, Wever-Pinzon O, Tracy CM, Bakhtina A, Bia R, Parnell T, Taleb I, et al. Distinct Transcriptomic and Proteomic Profile Specifies Patients Who Have Heart Failure With Potential of Myocardial Recovery on Mechanical Unloading and Circulatory Support. Circulation. 2022. doi: 10.1161/CIRCULATIONAHA.121.056600

34. Diakos NA, Navankasattusas S, Abel ED, Rutter J, McCreath L, Ferrin P, McKellar SH, Miller DV, Park SY, Richardson RS, et al. Evidence of Glycolysis Up-Regulation and Pyruvate Mitochondrial Oxidation Mismatch During Mechanical Unloading of the Failing Human Heart: Implications for Cardiac Reloading and Conditioning. JACC Basic Transl Sci. 2016;1:432–444. doi: 10.1016/j.jacbts.2016.06.009

35. Diakos NA, Selzman CH, Sachse FB, Stehlik J, Kfoury AG, Wever-Pinzon O, Catino A, Alharethi R, Reid BB, Miller DV, et al. Myocardial atrophy and chronic mechanical unloading of the failing human heart: implications for cardiac assist device-induced myocardial recovery. J Am Coll Cardiol. 2014;64:1602–1612. doi: 10.1016/j.jacc.2014.05.073

36. Szulik MW, Valdez S, Walsh M, Davis K, Bia R, Horiuchi E, O’Very S, Laxman AK, Sandaklie-Nicolova L, Eberhardt DR, et al. SMYD1a protects the heart from ischemic injury by regulating OPA1-mediated cristae remodeling and supercomplex formation. Basic Res Cardiol. 2023;118:20. doi: 10.1007/s00395-023-00991-6

37. Franklin S, Chen H, Mitchell-Jordan S, Ren S, Wang Y, Vondriska TM. Quantitative analysis of the chromatin proteome in disease reveals remodeling principles and identifies high mobility group protein B2 as a regulator of hypertrophic growth. Mol Cell Proteomics. 2012;11:M111 014258. doi: 10.1074/mcp.M111.014258

38. Mitchell-Jordan SA, Holopainen T, Ren S, Wang S, Warburton S, Zhang MJ, Alitalo K, Wang Y, Vondriska TM. Loss of Bmx nonreceptor tyrosine kinase prevents pressure overload-induced cardiac hypertrophy. Circ Res. 2008;103:1359–1362. doi: 10.1161/CIRCRESAHA.108.186577

39. Hickenlooper SM, Davis K, Szulik MW, Sheikh H, Miller M, Valdez S, Bia R, Franklin S. Correction to "Histone H4K20 Trimethylation Is Decreased in Murine Models of Heart Disease". ACS Omega. 2023;8:6124–6125. doi: 10.1021/acsomega.3c00112

40. Gibson-Corley KN, Olivier AK, Meyerholz DK. Principles for valid histopathologic scoring in research. Vet Pathol. 2013;50:1007–1015. doi: 10.1177/0300985813485099

41. Liu Y, Dostal DE, Tong CW. Isolation of Adult Mouse Cardiomyocytes Using Langendorff Perfusion Apparatus. Methods Mol Biol. 2021;2319:143–152. doi: 10.1007/978-1-0716-1480-8_16

42. Ye J, Coulouris G, Zaretskaya I, Cutcutache I, Rozen S, Madden TL. Primer-BLAST: a tool to design target-specific primers for polymerase chain reaction. BMC Bioinformatics. 2012;13:134. doi: 10.1186/1471-2105-13-134

43. Shibayama J, Yuzyuk TN, Cox J, Makaju A, Miller M, Lichter J, Li H, Leavy JD, Franklin S, Zaitsev AV. Metabolic remodeling in moderate synchronous versus dyssynchronous pacing-induced heart failure: integrated metabolomics and proteomics study. PLoS One. 2015;10:e0118974. doi: 10.1371/journal.pone.0118974

44. Kim MS, Pinto SM, Getnet D, Nirujogi RS, Manda SS, Chaerkady R, Madugundu AK, Kelkar DS, Isserlin R, Jain S, et al. A draft map of the human proteome. Nature. 2014;509:575–581. doi: 10.1038/nature13302

45. Tan X, Rotllant J, Li H, De Deyne P, Du SJ. SmyD1, a histone methyltransferase, is required for myofibril organization and muscle contraction in zebrafish embryos. Proc Natl Acad Sci U S A. 2006;103:2713–2718. doi: 10.1073/pnas.0509503103

46. Abu-Farha M, Lambert JP, Al-Madhoun AS, Elisma F, Skerjanc IS, Figeys D. The tale of two domains: proteomics and genomics analysis of SMYD2, a new histone methyltransferase. Mol Cell Proteomics. 2008;7:560–572. doi: 10.1074/mcp.M700271-MCP200

47. Hamamoto R, Furukawa Y, Morita M, Iimura Y, Silva FP, Li M, Yagyu R, Nakamura Y. SMYD3 encodes a histone methyltransferase involved in the proliferation of cancer cells. Nat Cell Biol. 2004;6:731–740. doi: 10.1038/ncb1151

48. Dreuw A, Radtke S, Pflanz S, Lippok BE, Heinrich PC, Hermanns HM. Characterization of the signaling capacities of the novel gp130-like cytokine receptor. J Biol Chem. 2004;279:36112–36120. doi: 10.1074/jbc.M401122200

49. Yang XO, Panopoulos AD, Nurieva R, Chang SH, Wang D, Watowich SS, Dong C. STAT3 regulates cytokine-mediated generation of inflammatory helper T cells. J Biol Chem. 2007;282:9358–9363. doi: 10.1074/jbc.C600321200

50. Yamamoto T, Sekine Y, Kashima K, Kubota A, Sato N, Aoki N, Matsuda T. The nuclear isoform of protein-tyrosine phosphatase TC-PTP regulates interleukin-6-mediated signaling pathway through STAT3 dephosphorylation. Biochem Biophys Res Commun. 2002;297:811–817. doi: 10.1016/s0006-291x(02)02291-x

51. Malagrino F, Puglisi E, Pagano L, Travaglini-Allocatelli C, Toto A. GRB2: A dynamic adaptor protein orchestrating cellular signaling in health and disease. Biochem Biophys Rep. 2024;39:101803. doi: 10.1016/j.bbrep.2024.101803

52. Wang D, Liu G, Meng Y, Chen H, Ye Z, Jing J. The Configuration of GRB2 in Protein Interaction and Signal Transduction. Biomolecules. 2024;14. doi: 10.3390/biom14030259

53. Wang J, Sun X, Wang X, Cui S, Liu R, Liu J, Fu B, Gong M, Wang C, Shi Y, et al. Grb2 Induces Cardiorenal Syndrome Type 3: Roles of IL-6, Cardiomyocyte Bioenergetics, and Akt/mTOR Pathway. Front Cell Dev Biol. 2021;9:630412. doi: 10.3389/fcell.2021.630412

54. Ikeda U, Ohkawa F, Seino Y, Yamamoto K, Hidaka Y, Kasahara T, Kawai T, Shimada K. Serum interleukin 6 levels become elevated in acute myocardial infarction. J Mol Cell Cardiol. 1992;24:579–584. doi: 10.1016/0022-2828(92)91042-4

55. Yamauchi-Takihara K, Ihara Y, Ogata A, Yoshizaki K, Azuma J, Kishimoto T. Hypoxic stress induces cardiac myocyte-derived interleukin-6. Circulation. 1995;91:1520–1524. doi: 10.1161/01.cir.91.5.1520

56. Tanaka T, Narazaki M, Kishimoto T. IL-6 in inflammation, immunity, and disease. Cold Spring Harb Perspect Biol. 2014;6:a016295. doi: 10.1101/cshperspect.a016295

57. Dawn B, Xuan YT, Guo Y, Rezazadeh A, Stein AB, Hunt G, Wu WJ, Tan W, Bolli R. IL-6 plays an obligatory role in late preconditioning via JAK-STAT signaling and upregulation of iNOS and COX-2. Cardiovasc Res. 2004;64:61–71. doi: 10.1016/j.cardiores.2004.05.011

58. Smart N, Mojet MH, Latchman DS, Marber MS, Duchen MR, Heads RJ. IL-6 induces PI 3-kinase and nitric oxide-dependent protection and preserves mitochondrial function in cardiomyocytes. Cardiovasc Res. 2006;69:164–177. doi: 10.1016/j.cardiores.2005.08.017

59. Eriksson U, Kurrer MO, Schmitz N, Marsch SC, Fontana A, Eugster HP, Kopf M. Interleukin-6-deficient mice resist development of autoimmune myocarditis associated with impaired upregulation of complement C3. Circulation. 2003;107:320–325. doi: 10.1161/01.cir.0000043802.38699.66

60. Prabhu SD. Cytokine-induced modulation of cardiac function. Circ Res. 2004;95:1140–1153. doi: 10.1161/01.RES.0000150734.79804.92

61. Yu X, Kennedy RH, Liu SJ. JAK2/STAT3, not ERK1/2, mediates interleukin-6-induced activation of inducible nitric-oxide synthase and decrease in contractility of adult ventricular myocytes. J Biol Chem. 2003;278:16304–16309. doi: 10.1074/jbc.M212321200

62. Hirota H, Yoshida K, Kishimoto T, Taga T. Continuous activation of gp130, a signal-transducing receptor component for interleukin 6-related cytokines, causes myocardial hypertrophy in mice. Proc Natl Acad Sci U S A. 1995;92:4862–4866. doi: 10.1073/pnas.92.11.4862

63. Chia YC, Kieneker LM, van Hassel G, Binnenmars SH, Nolte IM, van Zanden JJ, van der Meer P, Navis G, Voors AA, Bakker SJL, et al. Interleukin 6 and Development of Heart Failure With Preserved Ejection Fraction in the General Population. J Am Heart Assoc. 2021;10:e018549. doi: 10.1161/JAHA.120.018549

64. Diehl F, Brown MA, van Amerongen MJ, Novoyatleva T, Wietelmann A, Harriss J, Ferrazzi F, Bottger T, Harvey RP, Tucker PW, et al. Cardiac deletion of Smyd2 is dispensable for mouse heart development. PLoS One. 2010;5:e9748. doi: 10.1371/journal.pone.0009748

65. Yang D, Wei G, Long F, Nie H, Tian X, Qu L, Wang S, Li P, Qiu Y, Wang Y, et al. Histone methyltransferase Smyd3 is a new regulator for vascular senescence. Aging Cell. 2020;19:e13212. doi: 10.1111/acel.13212

66. Anderson DR, Poterucha JT, Mikuls TR, Duryee MJ, Garvin RP, Klassen LW, Shurmur SW, Thiele GM. IL-6 and its receptors in coronary artery disease and acute myocardial infarction. Cytokine. 2013;62:395–400. doi: 10.1016/j.cyto.2013.03.020

67. Markousis-Mavrogenis G, Tromp J, Ouwerkerk W, Devalaraja M, Anker SD, Cleland JG, Dickstein K, Filippatos GS, van der Harst P, Lang CC, et al. The clinical significance of interleukin-6 in heart failure: results from the BIOSTAT-CHF study. Eur J Heart Fail. 2019;21:965–973. doi: 10.1002/ejhf.1482

68. Frontiers Editorial O. An expression of concern on: Grb2 induces cardiorenal syndrome type 3: Roles of IL-6, cardiomyocyte bioenergetics, and Akt/mTOR pathway. Front Cell Dev Biol. 2022;10:1014332. doi: 10.3389/fcell.2022.1014332

69. Harhous Z, Booz GW, Ovize M, Bidaux G, Kurdi M. An Update on the Multifaceted Roles of STAT3 in the Heart. Front Cardiovasc Med. 2019;6:150. doi: 10.3389/fcvm.2019.00150

70. Jacoby JJ, Kalinowski A, Liu MG, Zhang SS, Gao Q, Chai GX, Ji L, Iwamoto Y, Li E, Schneider M, et al. Cardiomyocyte-restricted knockout of STAT3 results in higher sensitivity to inflammation, cardiac fibrosis, and heart failure with advanced age. Proc Natl Acad Sci U S A. 2003;100:12929–12934. doi: 10.1073/pnas.2134694100

71. Bourque G, Burns KH, Gehring M, Gorbunova V, Seluanov A, Hammell M, Imbeault M, Izsvak Z, Levin HL, Macfarlan TS, et al. Ten things you should know about transposable elements. Genome Biol. 2018;19:199. doi: 10.1186/s13059-018-1577-z

