## Supplementary figures and images for "Cardiomyocyte-specific loss of *Smyd5* leads to a robust activation of inflammatory signaling and heart failure in mice"

### Supplementary Figure S1

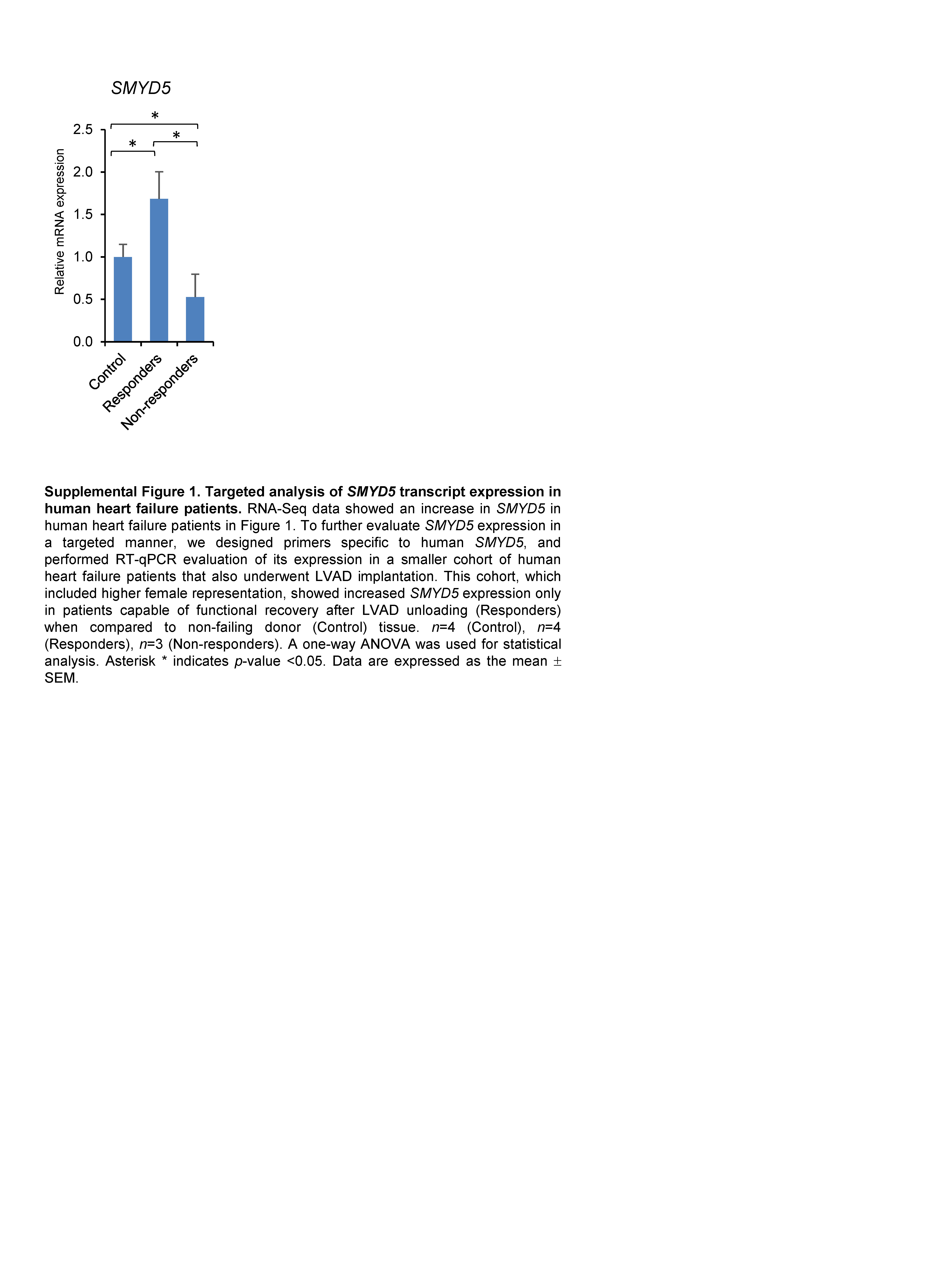

### Supplementary Figure S2

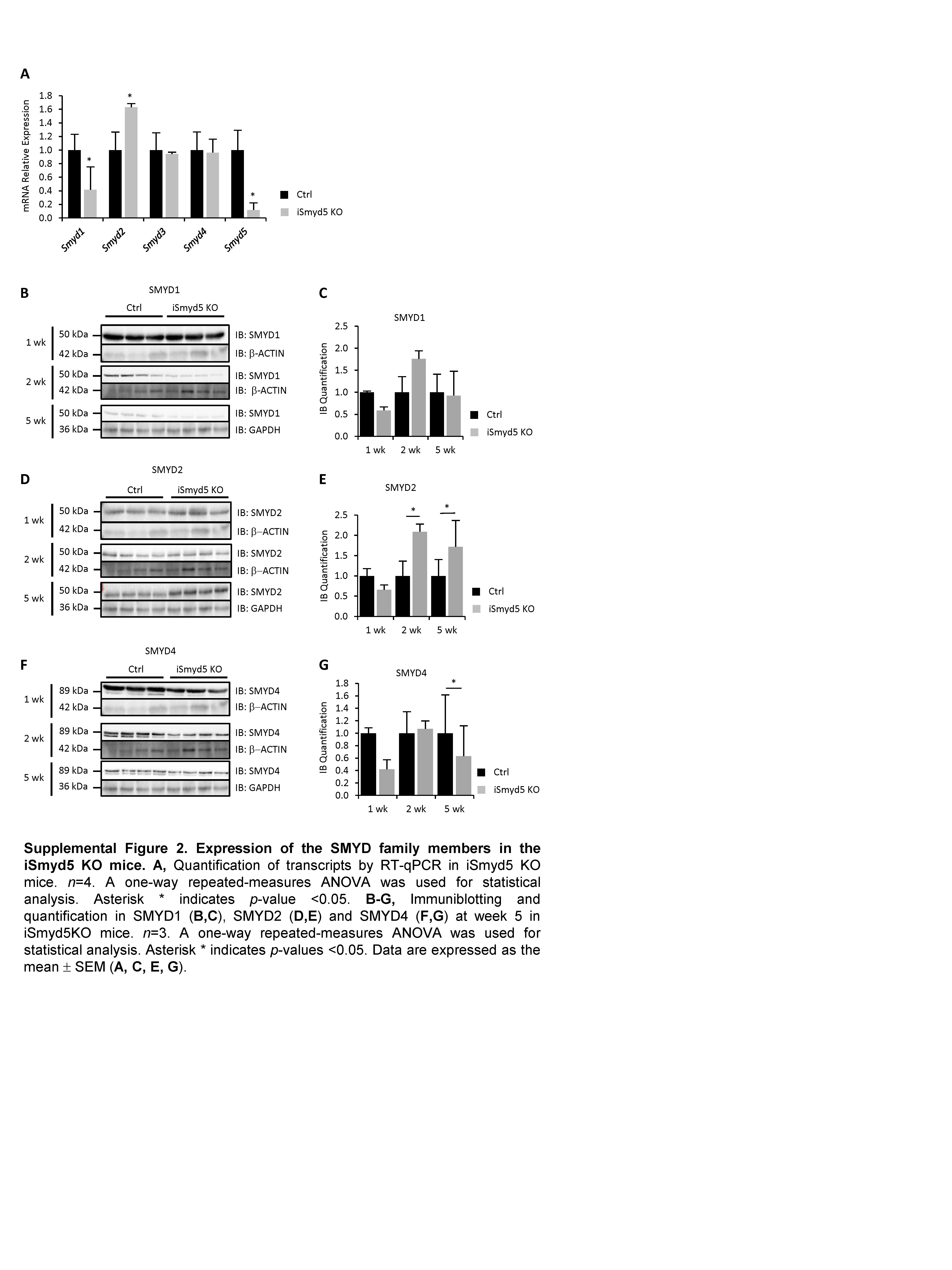

### Supplementary Figure S3

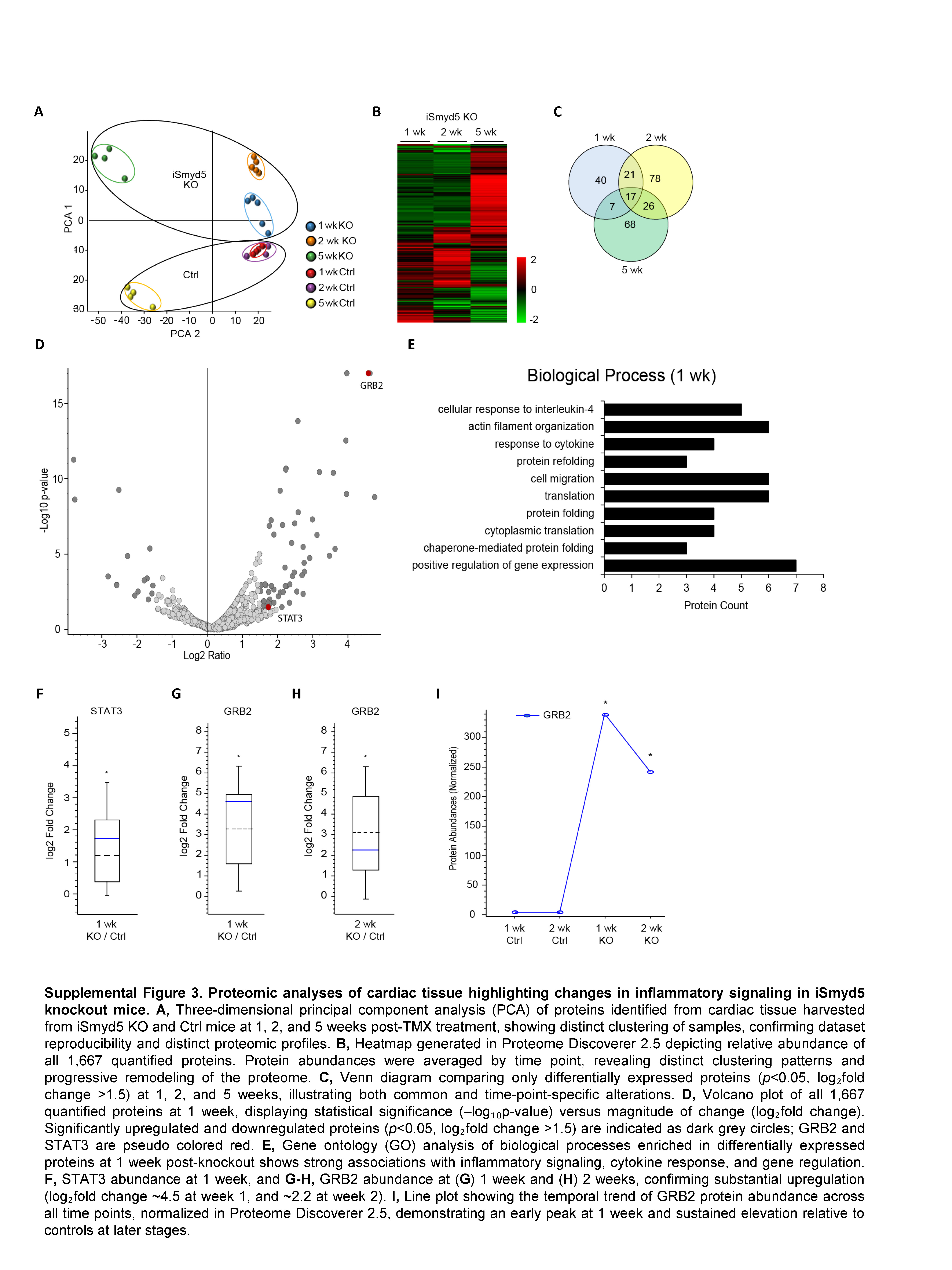
