## Supplementary Table S1 for "Cardiomyocyte-specific loss of *Smyd5* leads to a robust activation of inflammatory signaling and heart failure in mice"

**Table S1. Previously published studies on the histone methyltransferase SMYD5.** The data regarding SMYD5 is relatively sparse in the heart, but it has been previously studied in cultured cells, zebrafish, murine arthritis, and cancer, with 17 primary research articles focused on SMYD5 published to date.

| PMID | Publication Year | Authors | Title | Role | Journal/Book |
|---|---|---|---|---|---|
| 22921934 | 2012 | Stender JD, Pascual G, Liu W, Kaikkonen MU, Do K, Spann NJ, Boutros M, Perrimon N, Rosenfeld MG, Glass CK. | Control of proinflammatory gene programs by regulated trimethylation and demethylation of histone H4K20 | SMYD5 represses inflammatory cytokine gene transcription by depositing H4K20me3 at their promoters. | Mol Cell |
| 27377701 | 2016 | Fujii T, Tsunesumi S, Sagara H, Munakata M, Hisaki Y, Sekiya T, Furukawa Y, Sakamoto K, Watanabe S. | Smyd5 plays pivotal roles in both primitive and definitive hematopoiesis during zebrafish embryogenesis | SMYD5 is essential for hematopoiesis but not for muscle or heart development. | Sci Rep |
| 28250819 | 2017 | Kidder BL, Hu G, Cui K, Zhao K. | SMYD5 regulates H4K20me3-marked heterochromatin to safeguard ES cell self-renewal and prevent spurious differentiation | SMYD5 preserves heterochromatin and stem cell self-renewal by depositing H4K20me3. | Epigenetics Chromatin |
| 28951459 | 2017 | Kidder BL, He R, Wangsa D, Padilla-Nash HM, Bernardo MM, Sheng S, Ried T, Zhao K. | SMYD5 Controls Heterochromatin and Chromosome Integrity during Embryonic Stem Cell Differentiation | SMYD5 represses transposable elements and preserves genome integrity during differentiation. | Cancer Res |
| 33676231 | 2021 | Zhang Y, Hayden S, Spellmon N, Xue W, Martin K, Muzzarelli K, Kovari L, Yang Z. | Sperm chromatin-condensing protamine enhances SMYD5 thermal stability | Protamine binds SMYD5 and stabilizes it, indicating a potential role in spermatogenesis. | Biochem Biophys Res Commun |
| 35643234 | 2022 | Hou Y, Sun X, Gheinani PT, Guan X, Sharma S, Zhou Y, Jin C, Yang Z, Naren AP, Yin J, Denning TL, Gewirtz AT, Liu Y, Xie Z, Li C. | Epithelial SMYD5 Exaggerates IBD by Down-regulating Mitochondrial Functions via Post-Translational Control of PGC-1α Stability | SMYD5 methylates and degrades PGC-1α, impairing mitochondrial function and worsening IBD. | Cell Mol Gastroenterol Hepatol |
| 35182940 | 2022 | Aljazi MB, Gao Y, Wu Y, He J. | SMYD5 is a histone H3-specific methyltransferase mediating mono-methylation of histone H3 lysine 36 and 37 | SMYD5 mono-methylates H3K36 and H3K37, suggesting transcriptional regulatory activity. | Biochem Biophys Res Commun |
| 35218722 | 2022 | Chi G, Pei J, Li X, Li X, Pang H, Cui J, Wu D, Qu G, He Y. | SMYD5 acts as a potential biomarker for hepatocellular carcinoma | SMYD5 is overexpressed in HCC and promotes tumor proliferation and migration; a potential prognostic biomarker. | Exp Cell Res |
| 35740908 | 2022 | Zhang Y, Alshammari E, Sobota J, Yang A, Li C, Yang Z. | Unique SMYD5 Structure Revealed by AlphaFold Correlates with Its Functional Divergence | SMYD5 has a unique structure with an acidic cleft and lacks canonical methylation motifs, implying unique substrate targeting. | Biomolecules |
| 35680905 | 2022 | Zhang Y, Fang Y, Tang Y, Han S, Jia J, Wan X, Chen J, Yuan Y, Zhao B, Fang D. | SMYD5 catalyzes histone H3 lysine 36 trimethylation at promoters | SMYD5 catalyzes histone H3 lysine 36 trimethylation at promoters | Nat Commun |
| 36897778 | 2023 | Boehm D, Lam V, Schnolzer M, Ott M. | The lysine methyltransferase SMYD5 amplifies HIV-1 transcription and is post-transcriptionally upregulated by Tat and USP11 | SMYD5 enhances HIV transcription through interaction with Tat and methylation of Tat. | Cell Rep |
| 38723947 | 2024 | Tae IH, Ryu TY, Kang Y, Lee J, Kim K, Lee JM, Kim HW, Ko JH, Kim DS, Son MY, Cho HS. | Negative regulation of SH2B3 by SMYD5 controls epithelial-mesenchymal transition in lung cancer | SMYD5 represses SH2B3, enhancing EMT and metastatic potential. | Mol Cells |
| 39048817 | 2024 | Park J, Wu J, Szkop KJ, Jeong J, Jovanovic P, Husmann D, Flores NM, Francis JW, Chen YC, Benitez AM, Zahn E, Song S, Ajani JA, Wang L, Singh K, Larsson O, Garcia BA, Topisirovic I, Gozani O, Mazur PK. | SMYD5 methylation of rpL40 links ribosomal output to gastric cancer | SMYD5 methylates RPL40 to boost translation and tumor growth. | Nature |
| 39083378 | 2024 | Rafnsdottir S, Jang K, Halldorsdottir ST, Vinod M, Tomasdottir A, Möller K, Halldorsdottir K, Reynisdottir T, Atladottir LH, Allison KE, Ostacolo K, He J, Zhang L, Northington FJ, Magnusdottir E, Chavez-Valdez R, Anderson KJ, Bjornsson HT. | SMYD5 is a regulator of the mild hypothermia response | SMYD5 represses hypothermia-induced genes through H3K36me3; deletion enhances protective gene expression. | Cell Rep |
| 39103523 | 2024 | Miao B, Ge L, He C, Wang X, Wu J, Li X, Chen K, Wan J, Xing S, Ren L, Shi Z, Liu S, Hu Y, Chen J, Yu Y, Feng L, Flores NM, Liang Z, Xu X, Wang R, Zhou J, Fan J, Xiang B, Li E, Mao Y, Cheng J, Zhao K, Mazur PK, Cai J, Lan F. | SMYD5 is a ribosomal methyltransferase that catalyzes RPL40 lysine methylation to enhance translation output and promote hepatocellular carcinoma | SMYD5 methylates RPL40 K22 to enhance translation and support tumor progression. | Cell Res |
| 40165083 | 2025 | Xiao C, Su Z, Zhao J, Tan S, He M, Li Y, Liu J, Xu J, Hu Y, Li Z, Fan C, Liu X. | Novel regulation mechanism of histone methyltransferase SMYD5 in rheumatoid arthritis | SMYD5 implicated in inflammatory gene regulation contributing to RA pathogenesis. | Cell Mol Biol Lett |
| 40184250 | 2025 | Hamey JJ, Shah M, Wade JD, Bartolec TK, Wettenhall REH, Quinlan KGR, Williamson NA, Wilkins MR. | SMYD5 recognizes KXY motifs to trimethylate RPL40 K22, modulating translation. | SMYD5 represses inflammatory gene expression and is upregulated by LPS stimulation in macrophages. | Cell Rep |
